# Unbiased screen of RNA tailing enzymes at single-nucleotide resolution reveals a poly(UG) polymerase required for genome integrity and RNA silencing

**DOI:** 10.1101/422972

**Authors:** Melanie A. Preston, Douglas F. Porter, Fan Chen, Natascha Buter, Christopher P. Lapointe, Sunduz Keles, Judith Kimble, Marvin Wickens

## Abstract

Ribonucleotidyl transferases (rNTases) add non-templated ribonucleotides to diverse RNAs. We developed a screening strategy in *S. cerevisiae* to identify sequences added by candidate enzymes from different organisms at single-nucleotide resolution. The rNTase activities of 19 previously unexplored enzymes were determined. In addition to poly(A)- and poly(U)-adding enzymes, we identified a C-adding enzyme that is likely part of a two-enzyme system that adds CCA to tRNAs in a eukaryote; a nucleotidyl transferase that adds nucleotides to RNA without apparent nucleotide preference; and a poly(UG) polymerase, *C. elegans* MUT-2, which adds alternating U and G nucleotides to form poly(UG) tails. MUT-2 is known to be required for certain forms of RNA silencing, and mutations in the enzyme that are defective in silencing also fail to add poly(UG) tails in our assay. We propose that MUT-2 poly(UG) polymerase activity is required to promote genome integrity and RNA silencing.

## INTRODUCTION

Covalent modifications pervade biological regulation, and the discovery of enzymes that modify proteins and DNA have led to breakthroughs in metabolism, transcription and drug design. RNAs are extensively modified: 5’ termini are often capped, internal positions are altered both on ribose rings and bases, and 3’ termini receive untemplated nucleotides, referred to as “tails”. In eukaryotes, these 3’ tails control RNA stability, transport, processing and function, and affect virtually all classes of RNA, including mRNAs, snRNAs, tRNAs, lncRNAs and miRNAs. Tails, and the enzymes that add them, are critical in a wide spectrum of biological events. For example, uridylation is implicated in tumorigenesis, proliferation, stem cell maintenance, and the immune response^1-10^ and regulated poly(A) addition in early development, cancer, and memory^11-17^. Unbiased and global approaches with single-nucleotide resolution are needed to uncover new types of tails and alternate modification systems that may have gone unnoticed.

Members of the DNA polymerase β-like superfamily of nucleotidyl transferases catalyze non-templated addition of nucleotides^18,19^. Nucleotidyl transferases are related in amino acid sequence, but add nucleotides to divergent substrates, including RNAs, nucleotides, and antibiotics^19^. Nucleotidyl transferases that act on RNAs are referred to as ribonucleotidyl transferases (rNTases). rNTases include poly(A) polymerases (PAPs), poly(U) polymerases (PUPs; aka TUTases), and CCA-adding enzymes that add CCA tails to the 3’ end of tRNAs^20^. PAPs and PUPs cannot be distinguished unambiguously by inspection of their protein sequences.

Current methods to assay rNTase activity and nucleotide specificity generally are low-throughput and may not recapitulate rNTase specificities in living cells. *In vitro* approaches, which involve expression of recombinant protein or immunopurification, are dependent on assay conditions. Small molecules present *in vivo* can alter the specificities of tailing enzymes dramatically, complicating interpretation of *in vitro* studies^21^. Expression of candidate rNTases in *Xenopus* oocytes has enabled identification of multiple rNTases, but is low-throughput and not readily suitable for genome-wide analysis^16,22,23^.

We suspected that other tailing enzymes and forms of tails exist but have escaped detection. Powerful sequencing methods have been developed to identify tails on RNAs extracted from cells^24-26^. However, some tails may be added only at specific times or in certain cell types, occur on novel RNAs not commonly analyzed, or exist only transiently, perhaps triggering the RNA’s destruction. The challenge is to uncover all forms of tails, and identify the enzymes responsible, at a genome-wide scale.

We developed a screening approach to identify enzymes that add non-templated nucleotides to RNAs. Candidate rNTases were tethered *in vivo* to a reporter RNA in *S. cerevisiae*, and the number and identity of nucleotides they added were determined at single-nucleotide resolution using high-throughput sequencing. Our studies reveal previously undetected enzymes and tails, including a eukaryotic system with separate enzymes that add CC and A to form the ends of tRNAs, and a previously unknown enzymatic activity that adds alternating U and G residues to RNA 3’ termini. Mutations in the gene that encodes this poly(UG) polymerase are known to elevate transposition frequency^27-29^, disrupt silencing in the germline^30-34^, and impair RNA interference elicited by double-stranded RNA (RNAi)^35-39^. The same mutations abolish the enzyme’s poly(UG) addition activity. The poly(UG) polymerase, and likely poly(UG) tails, are required for these diverse RNA-dependent forms of regulation.

## RESULTS

### An *in vivo* tethering assay identifies rNTase activities

We devised an approach to classify rNTases by identification of the sequences they add to RNA (Fig. 1, Fig. S1). Candidate rNTases were fused to MS2 coat protein and an epitope tag (RGS-H_6_), and co-expressed in yeast with a reporter RNA that contained high-affinity MS2 binding sites. The interaction of MS2 coat protein with MS2 binding sites tethers the candidate protein to the reporter RNA, and circumvents RNA-binding proteins that might be required in a natural context^40^. We identified the nucleotides added by candidate enzymes by RT-PCR and high-throughput sequencing of whole-cell RNA extracted from yeast.

**Figure 1.**
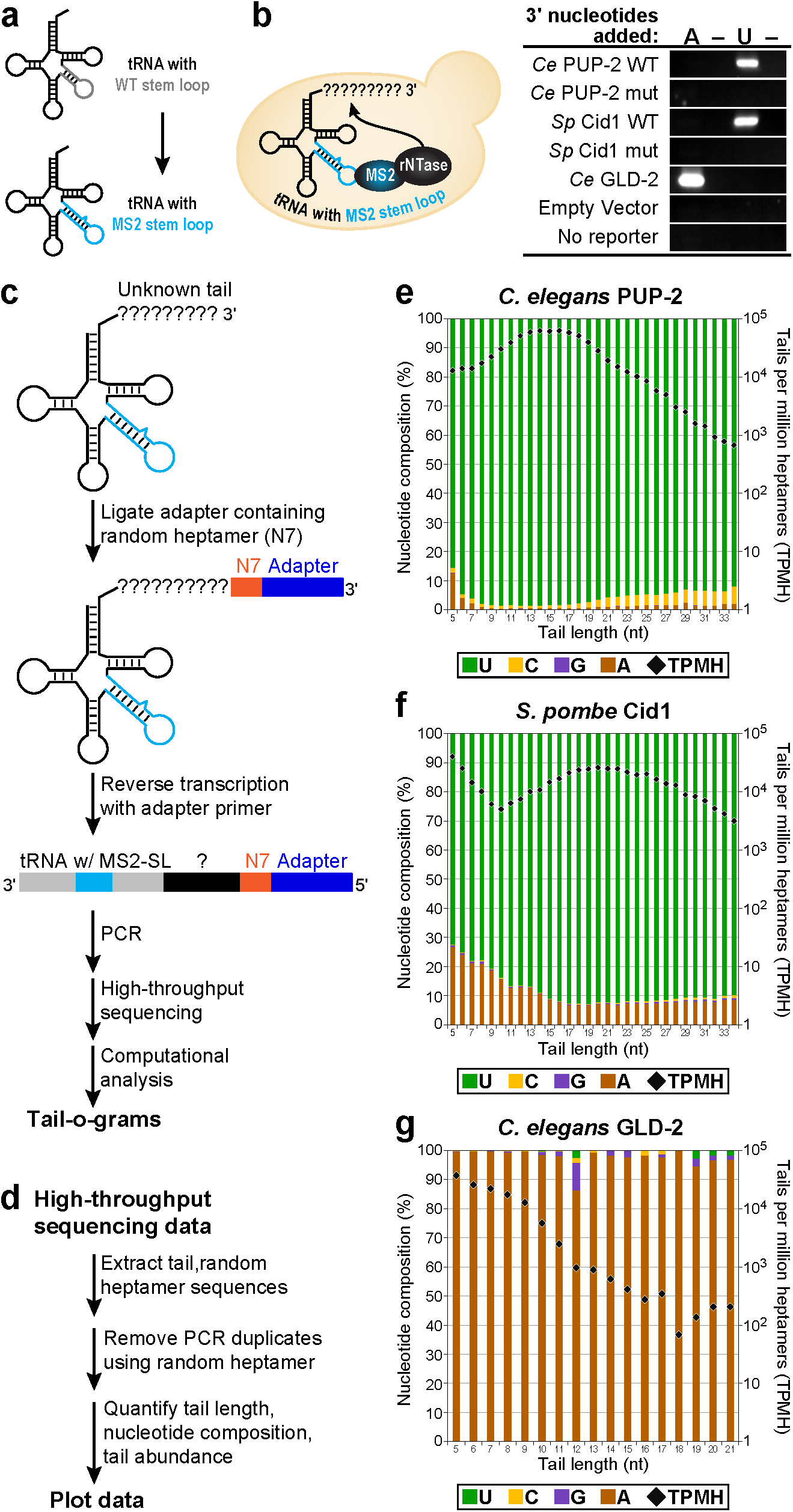
TRAID-Seq assay measures nucleotide addition activity *in vivo*. **(a, b)** TRAID-Seq strategy. **(a)** tRNA^Ser(AGA)^ variable arm (gray) is mutated to an MS2 stem loop (cyan) in the tRNA reporter. **(b)** Left, tRNA reporter is co-expressed with an MS2 coat protein-rNTase fusion in the yeast *S. cerevisiae*, and the tethered rNTase adds nucleotides to the 3’ end of the tRNA. Right, RT-PCR analysis to detect A tails or U tails added to the reporter tRNA by control rNTases, relative to empty vector or to a no-reporter control. Lanes marked with a dash indicate reactions performed without reverse transcriptase. **(c)** Schematic of sample processing. DNA adapter (dark blue) with a randomized nucleotide heptamer (N7, orange) is ligated to total RNA extracted from samples, followed by reverse transcription, PCR, and high-throughput sequencing**. (d)** Computational pipeline. **(e-g)** Tail-o-grams of nucleotides added by control rNTases. Data are shown for *C. elegans* PUP-2 (e), *S. pombe* Cid1 (f), and *C. elegans* GLD-2 (g). Percent of each nucleotide at each tail length is plotted on the y-axis. Each nucleotide is indicated by a different color: U (green), C (yellow), G (purple), A (brown). Percentages are given on the left y-axis. The number of tails detected per million heptamers (TPMH) are indicated by black diamonds, and correspond to the log scale on the right y-axis.

We identified an appropriate reporter RNA to serve as a substrate for rNTase enzymes. We first tested an RNase P-derived RNA that contained two MS2 binding sites^41,42^ (Fig. S1a,b). In cells expressing this RNA and an MS2 coat protein fusion with a known PUP (*C. elegans* PUP-2 or *S. pombe* Cid1), we detected addition of U tails to the reporter RNA, indicative of PUP activity (Fig. S1c); and this activity was not detected with a catalytically inactive form of the PUP. However, high background polyadenylation activity in yeast, observed in the absence of expressed rNTase enzymes, complicated analysis (Fig. S1c). We therefore created an alternative RNA substrate based on *S. cerevisiae* tRNA^Ser(AGA)^, a class II tRNA with a four base-pair variable arm. We replaced the variable arm with an MS2 binding site (Fig. 1a). Use of this substrate significantly reduced endogenous polyadenylation of the reporter RNA alone, as judged by reverse transcription and PCR (Fig. 1b), and enabled us to analyze MS2-rNTase fusion proteins unambiguously. We used this RNA in subsequent studies, and refer to it as “reporter tRNA” for simplicity.

Our approach accurately identified the activities of well-characterized rNTases. As proof-of-principle, we analyzed two known PUPs, *C. elegans* PUP-2^22^ (*Ce* PUP-2) and *S. pombe* Cid1^22,43^ (*Sp* Cid1), and a known PAP, *C. elegans* GLD-2^44^ (*Ce* GLD-2). The tails added *in vivo* to the reporter tRNA by each enzyme were analyzed using RT-PCR assays designed to detect U or A tails (Fig. 1a, Fig S1b). The U tail-specific primer yielded products with *Ce* PUP-2 and *Sp* Cid1 samples, while the A tail-specific primer yielded products only with *Ce* GLD-2. Tails were not detected when the active sites of *Ce* PUP-2 or *Sp* Cid1 were inactivated by point mutations (*Ce* PUP-2 mut, *Sp* Cid1 mut), nor when the reporter tRNA was expressed alone (Fig. 1b).

To identify tails of any nucleotide composition and length, we used high-throughput sequencing (Fig. 1c). Total RNA from each sample was ligated to a DNA adapter, such that the adapter was linked to the 3’ end of all RNAs in the sample. The presence of the adapter enabled detection of any nucleotides added and introduced a seven-nucleotide randomized sequence (random heptamer) that enabled us to remove PCR duplicates computationally. These features allowed us to analyze RNA molecules at single-nucleotide resolution. Following reverse transcription, samples were PCR-amplified using primers specific for the tRNA/MS2 stem loop and 3’ adapter sequences, and gel-purified products were subjected to paired-end sequencing on an Illumina platform.

To analyze the data, added tails first were extracted computationally, as defined by nucleotides between the 3’ end of the mature tRNA reporter (including the CCA sequence) and the random heptamer (Fig. 1d). We removed PCR duplicates and quantitated the number of unique tails and the composition of each nucleotide in the population of tails at each detected length (Fig 1d). Tail length, nucleotide composition, and the number of unique tails are plotted in “tail-o-grams” (Fig. 1e-g). In these plots, each tail length is assessed as a population to determine the percent of each nucleotide added among all tails of that length. Tails shorter than five nucleotides were discarded, as they were detected in the absence of the tethered enzymes and were random in sequence. To visualize the data, A, C, G and U are color-coded, and nucleotide compositions of tails are plotted relative to the length of tail added. Numbers of reads were normalized to the number of unique random heptamers (TPMH, tails per million heptamers) at each tail length, and displayed on a log scale.

The assay was accurate and sensitive, as judged by analyses of *Ce* PUP-2, *Sp* Cid1 and *Ce* GLD-2 enzymes. *Ce* PUP-2 and *Sp* Cid1 added tails primarily of uridines, and *Ce* GLD-2 added tails of adenosines (Fig. 1e-g), consistent with each of their known nucleotide specificities. Furthermore, the high sensitivity of the assay also enabled detection of secondary nucleotide addition preferences. For example, *Sp* Cid1 added uridine tails with 8.6% adenosine (Fig. 1f, Fig. 2c), consistent with its ability to add both A and U *in vitro*^43,45^.

**Figure 2.**
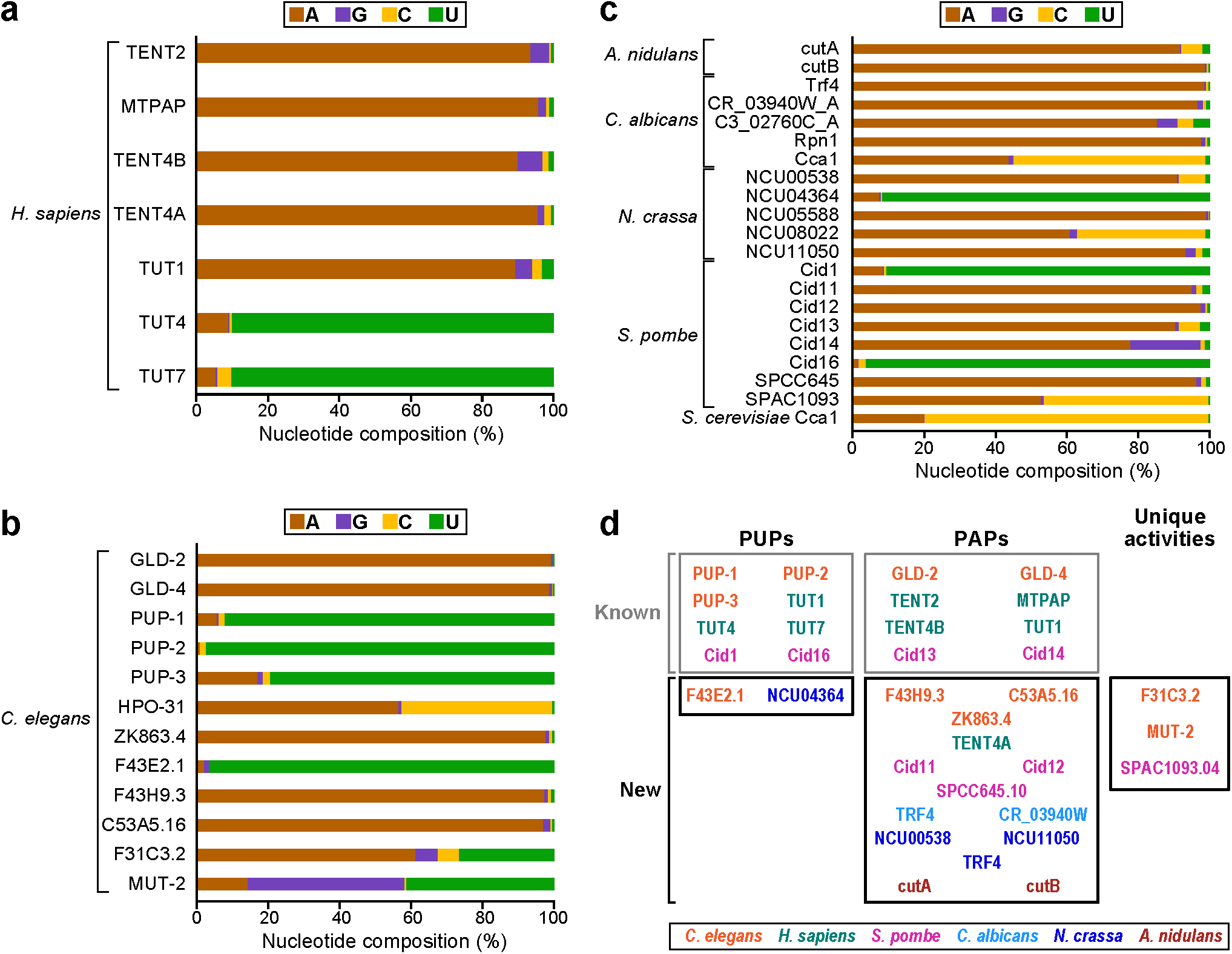
Analyses of nucleotide addition activities of 37 noncanonical rNTases from seven species. Overall percentages of each nucleotide added by **(a)** *H. sapiens*, **(b)** *C. elegans*, and **(c)** fungal rNTases. **(d)** Categorization of rNTases as PUPs, PAPs, or those of unique activity. rNTases are color-coded by organism. Gray boxes (top) indicate previously characterized (known) enzymes, and black boxes (bottom) indicate new enzymes.

Our assay, which we refer to as TRAID-Seq (tethered rNTase <u>a</u>ctivity <u>id</u>entified by high-throughput <u>seq</u>uencing), thus detects known rNTase activities. It circumvents the need for purified enzymes, which can be problematic with this class of proteins, and precisely identifies many thousands of independently captured tail sequences, enabling a sensitive determination of their sequences and relative abundances.

### New PUPs, PAPs, and CCA-adding enzymes

Using TRAID-Seq, we analyzed the nucleotide specificities of both characterized and previously untested rNTases. We tested 37 proteins from six species, including *Homo sapiens* (*Hs*), *Candida albicans* (*Ca*), *Neurospora crassa* (*Nc*), *Aspergillus nidulans* (*An*), *Schizosaccharomyces pombe* (*Sp*), and *Caenorhabditis elegans* (*Ce*) (Fig. 2, Table 1). Candidate rNTases were identified by the presence of a characteristic G(G/S) X_7-13_ DhDh motif and a downstream third aspartate^20^. To focus on noncanonical rNTases, we included putative rNTases with at least a partial type II nucleotide recognition motif (NRM)^19,20^, and excluded canonical rNTases, which are distinguished by the presence of a type I NRM^20^.

Nucleotide addition activities were classified first by the nucleotide composition of the tails added to the reporter tRNA (Fig. 2). For example, if tails added to the reporter tRNA consisted of primarily uridines, then the rNTase would be classified as a PUP. Through these analyses, we discovered 14 new PAPs and two new PUPs. We also identified likely CCA-adding enzymes in *N. crassa (Nc), C. albicans (Ca)* and *C. elegans (Ce)*, consistent with homology predictions in each respective curated database. These enzymes exhibit a preference for both C and A in the tails they add (Fig. 2b,c) and show an enrichment for the repeating CCA pattern within the tails added to the reporter tRNA. The p-values of CCA occurrence among the tails added by each enzyme, determined using a one-sided Wald’s test, are highly significant (adjusted p-values less than 1.6 x 10^-22^ (see “Supplemental Information: Methods”).

Enabled by the sensitivity of TRAID-Seq, we confirmed nucleotide specificities of previously characterized rNTases^16,21,22,43-53^ and identified surprising secondary preferences in certain enzymes. *Sp* Cid13 and *Sp* Cid14 are exemplary. Both were previously identified as PAPs^46^, yet both added other nucleotides as well. *Sp* Cid13 added 90.3% adenosine (s.d. 0.3%; n=4), and so was classified as a PAP, yet also added 6.0% cytosine (s.d. 0.3%; n=4; Fig. 2c, Fig. S2a). *Sp* Cid14 added 77.9% adenosine (s.d. 1.2%; n=3) and 19.7% guanosine (s.d. 0.8%; n=3; Fig. 2c, Fig. S2b). Analysis of the patterns of nucleotides added by enzymes with secondary preferences revealed no specific sequence motifs, in contrast to the enriched CCA pattern yielded by the CCA-adding enzymes.

In addition to identifying new PAPs, new PUPs, and CCA-adding enzymes (Fig. 2d), and discovering previously unappreciated nucleotide flexibilities, our analyses also revealed enzymes with previously undetected activities, as discussed in the following sections.

### C tails and a eukaryotic two-enzyme CCA-adding system

We identified an enzyme in *S. pombe* that primarily adds C nucleotides to RNAs. Based on sequence similarity, *S. pombe* SPAC1093.04 is predicted to be a CCA-adding enzyme, a highly conserved rNTase responsible for adding CCA to the 3’ end of virtually all tRNAs^54^. In TRAID-Seq with SPAC1093.04, we observed tails predominantly of oligo(C) or oligo(A) on reporter tRNAs with a CCA 3’ end (Fig. 2c; cytosine=46.0%, s.d. 6.0%; adenosine = 52.8%, s.d. 5.9%; n=5). Reporters with CC 3’ termini received almost exclusively oligo(C) (Fig. 3a, left, top). Tails added by *S. pombe* SPAC1093.04 and the *S. cerevisiae* CCA-adding enzyme (Cca1) clearly were distinct (Fig. 3a,b). The majority of tails added by *S. cerevisiae* Cca1 consist of repeating CCA motifs. In contrast, *Sp* SPAC1093.04 added long cytosine stretches (up to 19), which were often followed by a sequence of adenosines. The adenosines likely were added by endogenous PAPs in the TRAMP complex, which may recognize oligo(C)-tailed tRNAs as aberrant^47,50^. Differences between the activities of *Sp* SPAC1093.4 and *S. cerevisiae* Cca1 were manifest in computational analyses of sequence motifs of the tails they added. The trinucleotide CCA was highly enriched with the *S. cerevisiae* Cca1 but not *S. pombe* SPAC1093.04 (Fig. 3c, right). The products of both enzymes were significantly enriched for CC dinucleotides, as expected (Fig. 3c, left; for additional computational analyses, see Fig. S3). We conclude that *Sp* SPAC1093.04 possesses a distinctive C addition activity.

**Figure 3.**
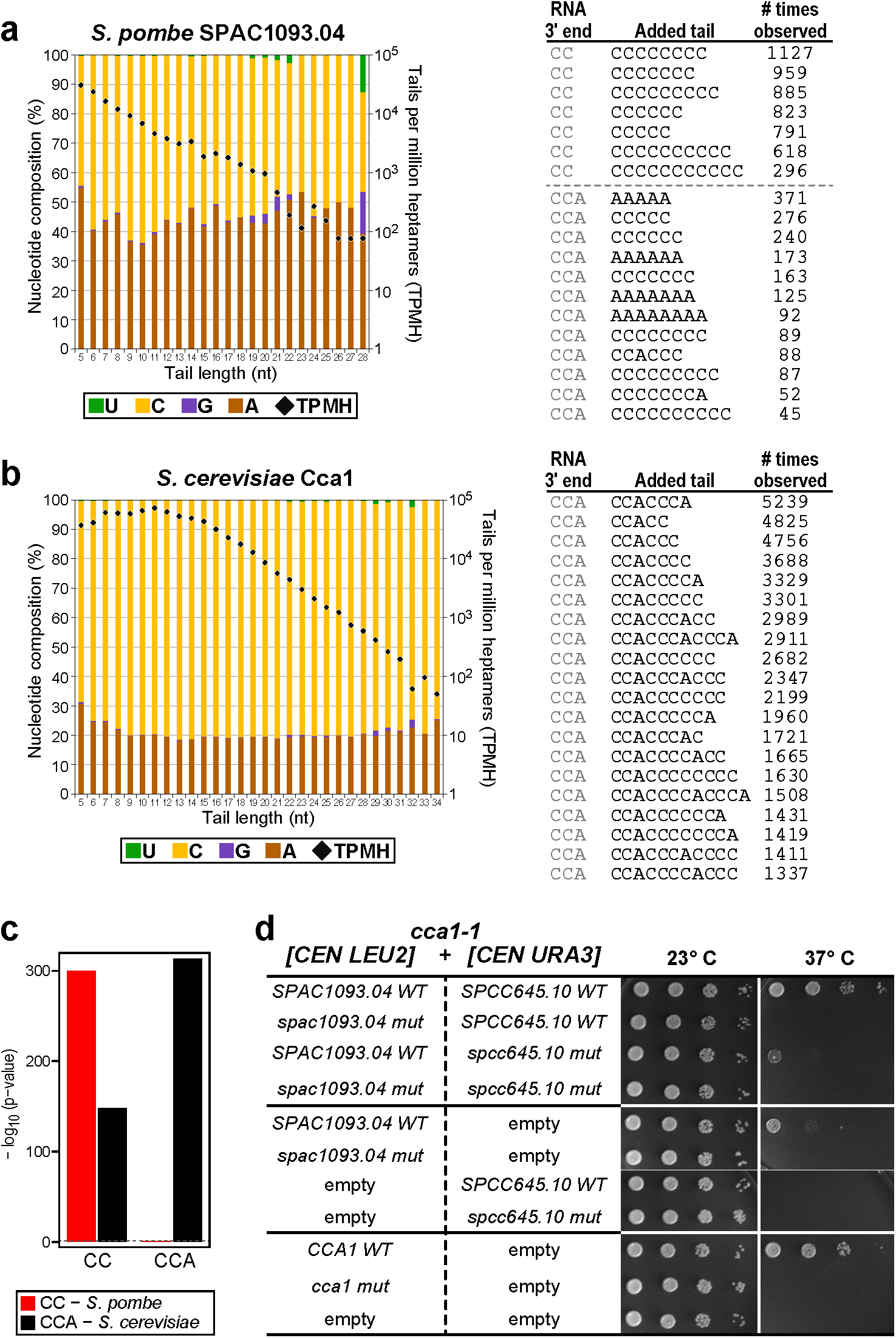
Nucleotide addition activity of *S. pombe* SPAC1093.04 and *S. cerevisiae* Cca1. **(a)** SPAC1093.04 adds tails enriched for polycytosine. Left, tail-o-gram depicting nucleotide composition in each added tail length and number of tails normalized to unique heptamer sequences. Right, most abundant tail sequences added to tRNA reporter containing a 3’ CC, or 3’ CCA end. **(b)** *S. cerevisiae* Cca1 adds cytosines and adenosines. Left, tail-o-gram depicting nucleotide composition in each added tail length and number of tails normalized to unique heptamer sequences. Right, most abundant tail sequences added to tRNA reporter containing a 3’ CCA end. **(c)** Sequence motif effect analysis of tails added by *Sp* SPAC1093.04 (red) and *Sc* Cca1 (black). Each adjusted p-value quantifies the significance of contribution of the indicated oligonucleotide to the variation in tail sequence read counts. Significances for dinucleotide (CC) and trinucleotides (CCA) after multiplicity correction with the Bonferroni procedure are shown. A dashed line indicates significance level 0.05. The −log_10_ p-values from left to right in the figure are 300, 148, 0.87, and 313. **(d)** Expression of both SPAC1093.04 and SPCC645.10, but of neither protein alone, rescues *cca1-1* temperature sensitivity. *cca1-1* mutant strains containing *CEN* plasmids expressing indicated plasmids were serially diluted, spotted on SD-Ura-Leu media and grown at 37°C for 3 days or 23°C for 4 days.

The *S. pombe* genome encodes a second enzyme (SPCC645.10) with sequence similarity to CCA-adding enzymes. This enzyme yielded tails of almost entirely adenosines (Fig. 2c, 96.3%, s.d. 0.7%). Thus, we wondered whether *Sp* SPAC1093.04 and *Sp* SPCC645.04 might act sequentially to add CCA to tRNAs, with *Sp* SPAC1093.04 first adding two C’s and then *Sp* SPCC645.10 adding the terminal A. The use of two enzymes to add CCA has not previously been demonstrated in eukaryotes, though it occurs in certain bacteria^55,56^.

To test our hypothesis, we determined whether the two *S. pombe genes*, SPAC1093.04 and SPCC645.04, could rescue lethality due to loss of CCA-adding activity in *S. cerevisiae*. We used a *cca1-1* mutant strain of *S. cerevisiae* strain bearing a temperature-sensitive (*ts*) allele of the essential *CCA1* gene. *CCA1* encodes the single protein that adds CCA to tRNAs in *S. cerevisiae*^57,58^. SPAC1093.04 and SPCC645.10 were expressed in the *cca1-1* strain using the *CCA1* promoter and terminator sequences on single-copy plasmids. Effects on temperature sensitivity were assessed in strains expressing the *S. pombe* proteins either together or with an empty vector (Fig. 3d, Fig. S4).

Coexpression of both *S. pombe* enzymes rescued loss of endogenous CCA addition activity in *S cerevisiae*. *cca1-1* temperature sensitivity at 37°C was fully rescued by co-expression of SPAC1093.04 and SPCC645.10, and by the wild-type *CCA1* positive control. Expression of SPAC1093.04 alone only partially suppressed the *cca1-1 ts* phenotype. Expression of SPCC645.10 alone or catalytic-inactive versions of SPAC1093.04 and SPCC645.10 failed to rescue the temperature sensitivity. Thus, our data suggest that SPAC1093.04 and SPCC645.10 collaborate to add CCA to tRNAs to rescue the *cca1-1 ts* phenotype. We propose that this collaboration is also necessary for CCA addition to tRNAs in *S. pombe* because both enzymes are essential^59,60^. To our knowledge, this would be the first dual enzyme system that adds CCA to tRNAs in a eukaryote.

### An enzyme with broad specificity

*C. elegans* F31C3.2 displayed a uniquely broad nucleotide specificity (Fig. 2b, Fig. 4a). The majority of nucleotides added by the enzyme were adenosines and uridines, but guanosines and cytosines were also prominent (Fig. 4b). Analysis of enrichment of short oligonucleotide sequences within the added tails yielded no discernible pattern or sequence motif (of p-value less than 0.05). This is further emphasized by computational analysis of all 16 possible dinucleotide sequences, none of which displays statistically significant enrichment among the added tails (Fig. S5). The base composition of the added tails paralleled intracellular ribonucleotide concentrations in *S. cerevisiae*^61^ (Fig. 4b). Taken together with the random nature of the sequences added, we suggest that *Ce* F31C3.2 may be relatively indiscriminate in its nucleotide preference. Hereafter, we refer to *Ce* F31C3.2 as nucleotide polymerase-1 (NPOL-1).

**Figure 4.**
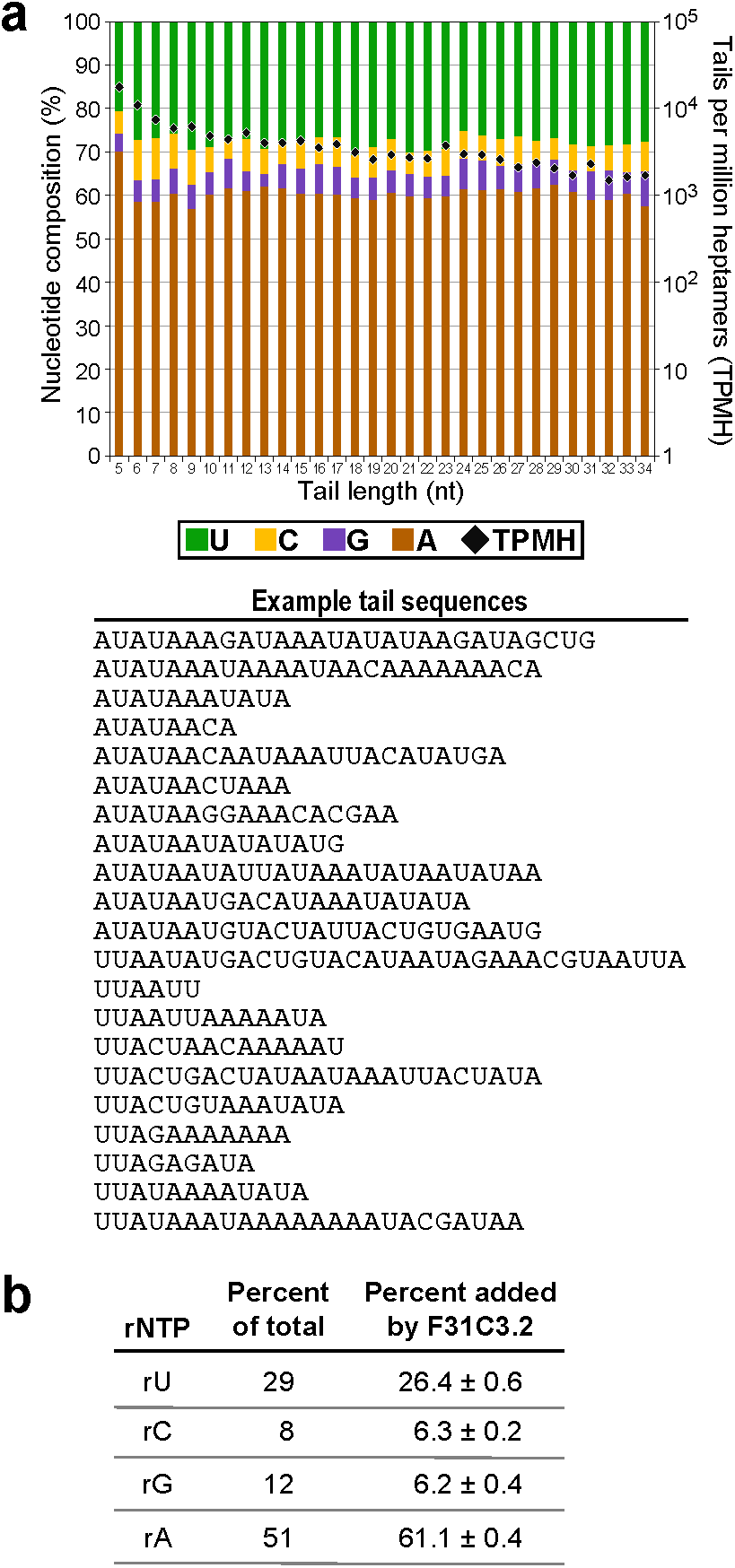
*Ce* F31C3.2 (NPOL-1) adds nucleotides without sequence motifs. **(a)** Top, tail-o-gram of *Ce* F31C3.2 activity. Bottom, example tail sequences added by *Ce* F31C3.2. **(b)** *Ce* F31C3.2 addition vs. measured rNTP concentrations in yeast^61^.

### A poly(UG) polymerase required for RNA silencing

*C. elegans* MUT-2 protein yielded tails with a 1:1 ratio of uridines and guanosines (Fig. 2b, Fig. 5a). Surprisingly, we found that *Ce* MUT-2 added alternating U and G nucleotides, yielding striking, polymeric sequences of alternating U and G (Fig. 5b). Computational analysis confirmed repetitive UG addition, and revealed that tails began with either uridine or guanosine. Of the two predicted splicing isoforms of *Ce* MUT-2 (mut-2a, mut-2b, https://wormbase.org/species/c_elegans/gene/WBGene00003499#0-1-3), only MUT-2a protein exhibited polymerase activity (Fig. 6b). We refer to this enzyme as a poly(UG) polymerase.

**Figure 5.**
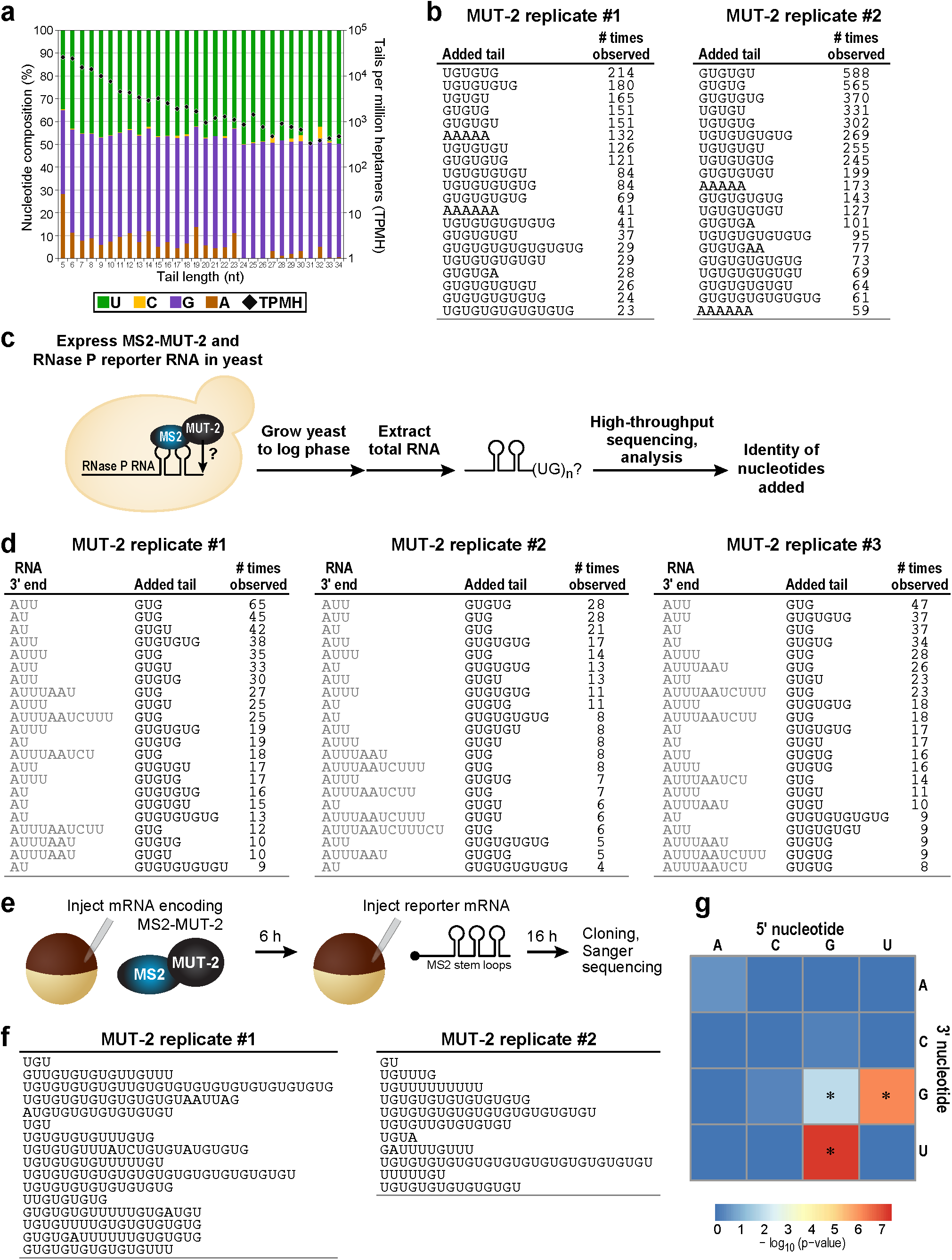
*Ce* MUT-2 is a poly(UG) polymerase. **(a)** Tail-o-gram depicting *Ce* MUT-2 nucleotide addition activity in yeast. **(b)** The most abundant tail sequences identified in two biological replicates of *Ce* MUT-2 TRAID-Seq assays. **(c)** Schematic of experiment to test *Ce* MUT-2 activity on RNase P RNA reporter in yeast. **(d)** *Ce* MUT-2 adds UG repeats to RNase P RNA in three biological replicates, regardless of the 3’ end sequence of the reporter RNA. **(e)** Schematic of experiment to test *Ce* MUT-2 activity in *Xenopus laevis* oocytes. **(f)** UG tail sequences from two biological replicates of *Ce* MUT-2 activity in *Xenopus laevis* oocytes. From replicate 1, we cloned 43 independent reporter sequences, 16 had added tails, and all contained UG. From replicate 2, we cloned 31 independent reporter sequences, 11 had added tails, and all contained UG. **(g)** Statistical analysis of all possible dinucleotides in the tails added by MUT-2. A heatmap of p-values for individual dinucleotides with minus logarithm (base 10) is shown. Each p-value quantifies the significance of adjusted contribution of each dinucleotide to the variation in tail sequence read counts. Dinucleotides with a significant effect after multiplicity correction at significance level 0.05 are marked with an asterisk (*).

**Figure 6.**
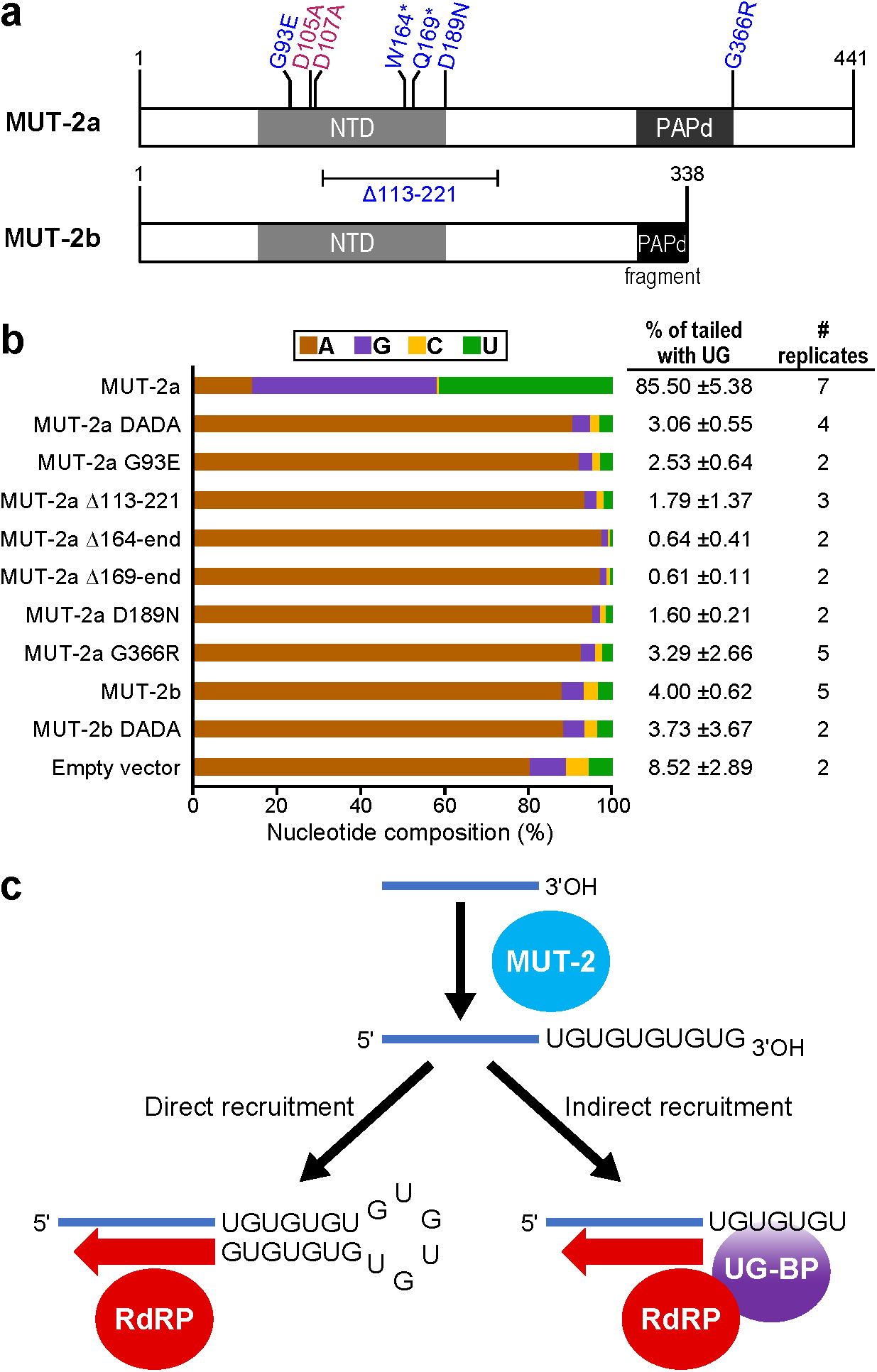
*Ce* MUT-2 mutants defective for RNAi lack poly(UG) polymerase activity. **(a)** Schematic of *Ce* MUT-2 isoforms and tested mutations, known catalytic mutants (pink), mutants identified in forward genetic screen^35^ (blue). NTD, Nucleotidyl transferase domain; PAPd, Poly(A) polymerase-associated domain. **(b)** Percent of nucleotides added by each *Ce* MUT-2 enzyme variant. Percent of tails containing UG repeats, standard deviation, and number of biological replicates are indicated. **(c)** Model depicting the potential roles of poly(UG) tails in small RNA amplification in *C. elegans*. Poly(UG) tails could directly recruit RNA-dependent RNA polymerase (RdRP) for small RNA amplification (left). Alternatively, poly(UG) tails could be identified by a poly(UG) binding protein (UG-BP), which then recruits an RdRP (right).

**Table 1:**

| rNTase | Species | Characterization | Nucleotide Preference | Histidine in NRM |
| --- | --- | --- | --- | --- |
| Cid1 | *S. pombe* | Kwak & Wickens 2007;<br>Rissland *et al.* 2007 | U | yes |
| PUP-1 | *C. elegans* | Kwak & Wickens 2007 | U | yes |
| PUP-2 | *C. elegans* | Kwak & Wickens 2007 | U | yes |
| TUT4 | *H. sapiens* | Kwak & Wickens 2007;<br>Rissland *et al.* 2007 | U | yes |
| TUT7 | *H. sapiens* | Kwak & Wickens 2007;<br>Rissland *et al.* 2007 | U | yes |
| NCU04364.7 | *N. crassa* | This study | U | yes |
| **Cid16** | ***S. pombe*** | **This study** | **U** | **no (Lys)** |
| **PUP-3** | ***C. elegans*** | **Kwak & Wickens 2007** | **U** | **no (Arg)** |
| **F43E2.1** | ***C. elegans*** | **This study** | **U** | **no (Arg)** |
| **F31C3.2** | ***C. elegans*** | **This study** | **A, U, C, G**<br>**(indiscriminate)** | **yes** |
| **MUT-2** | ***C. elegans*** | **This study** | **U,G** | **yes** |
| **TUT1**<br>**(Star-PAP)** | ***H. sapiens*** | **Trippe *et al.* 2006;**<br>**Mellman *et al.* 2008** | **A** | **yes** |
| cutB | *A. nidulans* | This study | A | no (Asn) |
| Trf4 | *S. cerevisiae* | Kadaba *et al.* 2004 | A | no (Asn) |
| Trf5 | *S. cerevisiae* | Haracska *et al.* 2005 | A | no (Asn) |
| Trf4 | *C. albicans* | This study | A | no (Asn) |
| CR_03940W_A | *C. albicans* | This study | A | no (Asn) |
| GLD-2 | *C. elegans* | Wang *et al.* 2002 | A | no (Asn) |
| GLD-4 | *C. elegans* | Schmid *et al.* 2009 | A | no (Asn) |
| TENT2 (GLD2) | *H. sapiens* | Kwak *et al.* 2004 | A | no (Asn) |
| TENT4B (PAPD5) | *H. sapiens* | Rammelt *et al.* 2011 | A | no (Asn) |
| TENT4A | *H. sapiens* | This study | A | no (Asn) |
| TRF4 | *N. crassa* | This study | A | no (Asn) |
| Cid12 | *S. pombe* | This study | A | no (Asn) |
| Cid14 | *S. pombe* | Read *et al.* 2002 | A | no (Asn) |
| cutA | *A. nidulans* | This study | A | no (Arg) |
| NCU00538.7 | *N. crassa* | This study | A | no (Arg) |
| Cid11 | *S. pombe* | This study | A | no (Arg) |
| Cid13 | *S. pombe* | Read *et al.* 2002;<br>Saitoh *et al.* 2002 | A | no (Arg) |
| MTPAP | *H. sapiens* | Tomecki *et al.* 2004 | A | no (Leu) |
| NCU11050.7 | *N. crassa* | This study | A | no (Leu) |
| C53A5.16 | *C. elegans* | This study | A | no (Pro) |
| F43H9.3 | *C. elegans* | This study | A | no (not aligned) |
| RPN1 | *C. albicans* | This study | A | no (Glu) |

To test whether this unusual specificity was independent of the RNA substrate, we used a different RNA, derived from RNase P RNA (Fig. 5c, Fig. S1). This RNA neither had a CCA 3’ end nor resembled a tRNA. *Ce* MUT-2 again added tandem UG repeats, as demonstrated by representative sequences from three biological replicates (Fig. 5d). Indeed, it added alternating UG to any of the multiple termini formed on the RNase P reporter RNA.

To further examine whether addition of UG repeats was intrinsic to the protein, we tested *Ce* MUT-2 in a different organism and cell type – *Xenopus laevis* oocytes. *Ce* MUT-2 was expressed in *X. laevis* oocytes by microinjecting an mRNA encoding *Ce* MUT-2a fused to MS2 coat protein. After allowing the protein to accumulate, an RNA containing a polylinker and three MS2 loops was injected (plgMS2-luc, Fig. 5e). Untemplated nucleotides were detected on 35-37% of the reporter RNA molecules, and all of these tails contained UG repeats, most commonly tandem UG repeats (Fig. 5f). We also observed a few instances of short uridine stretches, perhaps due to endogenous *Xenopus* TUT4 or TUT7 poly(U) polymerase activity on reporter RNAs containing UG repeats.

To evaluate the data statistically, we compiled the sequences of *Ce* MUT-2-catalyzed tails from all TRAID-Seq experiments and examined the statistical significance of the occurrence of each of the possible 16 dinucleotide pairs (Fig. 5g). 5’ -GU-3’ and 5’ -UG-3’ were highly enriched, with −log_10_ (p-values) of 7.3 and 6.2. We conclude that *Ce* MUT-2 catalyzes the addition of alternating UG. To our knowledge, *Ce* MUT-2 is the first example of a poly(UG) polymerase. The UG repeats are essentially perfectly repeated throughout the tails added, a remarkable pattern not observed in sequences added by other known nucleotidyl transferases.

The UG-adding activity of *Ce* MUT-2 likely is critical for RNAi. *Ce* MUT-2 was first identified in a screen for mutants with increased frequency of transposon Tc1 excision in the *C. elegans* germline^27^. The same gene was later identified in a forward genetic screen designed to detect genes involved in the efficacy of “feeding RNAi” in *C. elegans,* and so was referred to as RDE-3 (“RNAi-defective”)^35^. *Ce* MUT-2 has since been implicated in the production of secondary small RNAs (22G RNAs)^35,37^. The original RNAi-defective screen yielded six independent alleles, each of which alters a region predicted to be important for catalytic activity (Fig. 6a). We tested the nucleotide addition activities of MUT-2 fusion proteins that correspond to each of these mutants, as well as a negative control D105A/D107A (DADA) in which two predicted catalytic aspartates were mutated to alanine (Fig. 6b). All of the *Ce* MUT-2 mutations identified previously^35^ lacked UG addition activity, and the nucleotide compositions of any tails added resembled the negative control and the catalytically inactive enzyme (DADA mutant). Because *C. elegans* mutant strains harboring these same alleles are defective for multiple forms of RNA silencing and for secondary small RNA production in RNAi interference, we propose that poly(UG) polymerase activity is important in those events.

## DISCUSSION

TRAID-Seq is facile, sensitive, and enables parallel analyses of many rNTases. Thousands of independently captured tail sequences are identified, permitting sensitive determinations of their sequences and abundances, and thus the identity and patterns of nucleotides added. Since neither the protein nor its RNA product need to be purified, multiple proteins can be analyzed in parallel and thousands of tail sequences determined precisely. While we tested proteins identified through their sequence similarity to rNTases, the approach could be applied to a much broader range of ORFs, and identify enzymes that catalyze any RNA modifications detectable through sequencing, including certain base modifications.

Our analyses reveal the activities of 19 previously uncharacterized members of the rNTase protein family, from six species. The active site regions of the PAPs and PUPs we identified bear on how U and A are distinguished by different enzymes in the same family. Prior work demonstrated that a histidine in the active-site regions of Human Gld2 and *S. pombe* Cid1 dictates their apparent preferences for A and U, respectively^62-67^. Similarly, U-adding enzymes appear to have arisen repeatedly in evolution by the insertion of histidine into ancestral A-adding enzymes^62^. However, among the U-adding enzymes we uncovered here, several (*Sp* Cid16, *Ce* PUP-3 and *Ce* F43E2.1) lack that histidine, and one that possesses a histidine (*Hs* TUT1/Star-PAP) adds adenosines^21^ (Table 1). Both *Ce* NPOL-1, the broad specificity NTase, and *Ce* MUT-2, which adds alternating U and G, possess a histidine, further emphasizing that purines can be accommodated. These findings illustrate that the basis of nucleotide discrimination is more complex than previously thought. Analysis of the structures of these enzymes bound to their nucleotide substrates should be illuminating.

Our findings suggest that protein partners or small molecules may contribute to the specificities of certain nucleotidyl transferases. *In vivo, Hs* TUT1 (also known as Star-PAP) adds U’s to U6-snRNA^51^, but adds A’s to a variety of mRNAs^21,68,69^. In TRAID-Seq, we detected a strong preference for A (adenosine= 89.5%, s.d. 1.4%), and only low levels of incorporation of other nucleotides (uridine=3.2%, s.d. 0.7%; cytosine= 2.5%, s.d. 0.4%; guanosine=4.8%, s.d. 0.6%). A specific phosphoinositide enhances A addition activity of this enzyme *in vitro*^21^, and may underlie these differences. *Aspergillus (An)* CutA adds CU-rich 3’ terminal extensions to RNAs *in vivo* and prefers CTP *in vitro*^70,71^. In TRAID-Seq, *An* CutA added predominantly adenosine (91.8%, s.d. 0.4%) vs C (5.9%, s.d. 0.2%) or U (1.9%, s.d. 0.3%). *In vivo* in Aspergillus, *An* CutB collaborates with *An* CutA to form CU-rich tails^72^ but added virtually all A’s in TRAID-Seq [98.7%, s.d. 0.2% vs. C (0.4%, s.d. 0.05%) or U (0.3%, s.d. 0.07%)]. These findings suggest that additional cofactors or the nature of the RNA substrates can influence specificity *in vivo*.

The sensitivity of TRAID-Seq revealed previously undetected nucleotide addition capabilities that may underlie the addition of *in vivo* tails that have been enigmatic. For example, three human PAPs (TENT2, TENT4b, and TUT1) are capable of G addition, albeit at a low level in our system (Fig. 2a), and could explain the observation of G addition on mRNAs in human cells^24^. Indeed, TENT4a and TENT4b were recently implicated in G addition to mRNAs, which then are protected from deadenylation^73^. Perhaps the ability of several human PAPs to add G’s might indicate that other classes of RNAs are subject to such regulation. The abilities of *Sp* Cid13 and *Sp* Cid14 to add C and G, respectively, in addition to A, and suggests an analogous mechanism of RNA regulation in *S. pombe*.

*C. elegans* NPOL-1 added tails composed of all four nucleotides without a discernible sequence pattern, and is distinct in specificity from the other enzymes tested. The levels of incorporation mirror intracellular concentrations of ribonucleoside triphosphate concentrations, which may determine the proportions of nucleotides added. The broad specificity of *Ce* NPOL-1 echoes the activities of terminal deoxynucleotidyl transferase (TdT) and *E. coli* poly(A) polymerase (EcPAP), which also can add all four nucleotides without a template^74-76^. However, NPOL-1 diverges in sequence from TdT and EcPAP, which belong to a different subfamily of nucleotidyl transferases^19^. Indeed, the closest ortholog of NPOL-1 is human TENT2 (aka GLD2/PAPD4; 37% sequence homology; https://wormbase.org/species/c_elegans/gene/WBGene00001596#0-1-3). The addition of random nucleotides within, or at the end of, homopolymeric tails could interfere with their function^26,73,77^. It will be of interest to test whether binding partners, RNA substrates, or cofactors alter the nucleotide preferences of NPOL-1, as has been observed with *Hs* TUT1/Star-PAP^69^.

We propose that SPAC1093.04 and SPCC645.10 constitute a two-enzyme system that catalyzes CCA addition to tRNAs in *S. pombe*. This is strongly suggested by their specificities and ability to jointly complement a *S. cerevisiae* strain lacking a functional CCA-adding enzyme. This would be the first report of a two-enzyme CCA addition system in a eukaryote. Studies in *S. pombe* will test this proposal in its natural context, and determine whether either these enzymes act on other RNAs as well.

MUT-2, the poly(UG) polymerase, is remarkable both in its enzyme activity and roles in RNA biology. Its capacity to polymerize tails composed of as many as 18 perfect UG repeats is striking. Even longer UG tails likely were present, but were undetected due to sequencing read limitations. Alternating U and G addition bears comparison to that of CCA-adding enzymes, which switch nucleotide specificities as they sequentially add C, C and then A to a tRNA. They do so through a single active site, repositioning the growing 3’ end relative to the enzyme^78-82^. Redesign of the polarity of hydrogen bonds in a CCA-adding enzyme enable it to add UUG to a tRNA substrate in vitro^76^ and two CCAs can be added by shifting the 3’ end relative to the protein^83,84^. Repetitive UG addition by *Ce* MUT-2 may be promoted by repositioning the 3’ -most UG relative to the *Ce* MUT-2 active site.

The functions of *Ce* MUT-2 *in vivo* are diverse. *mut-2* was first isolated in a genetic screen for elevated transposition frequency in *C. elegans*^27^, and later in a screen for mutants with impaired RNAi in response to exogenous double-stranded RNA^35^. *mut-2* mutants possess reduced levels of secondary small RNAs^35,37,39^ (22G and 26G RNAs), suggesting that the protein stabilizes or helps to generate them. *Ce* MUT-2 function *in vivo* likely hinges on its poly(UG) polymerase activity, since the mutations identified in RNAi-defective *mut-2* mutants abrogate poly(UG) polymerase activity in our assays (Fig. 6a,b).

The multiple roles of *Ce* MUT-2 – preserving genome integrity^27-29^, silencing transgenes^30-34^ and promoting RNAi due to exogenous dsRNA^35-39^ – all likely reflect the same underlying molecular mechanisms. MUT-2 increases the abundance of secondary RNAs during RNAi, suggesting that UG tails are important in RdRP-based secondary siRNA synthesis or stabilization^35,37^. In one simple model, MUT-2 adds poly(UG) to the 3’ end of sliced RNAs generated in an Ago-dependent process. The poly(UG) tails would then provide a distinctive mark on sliced RNAs and bind RdRP directly, or via a separate UG-binding protein (Fig 6c). In either case, the tail could be single-stranded, or, as we favor, form a more complex structure involving U-G, U-U, or G-G pairing interactions (depicted as UG pairing in Fig. 6c, left). By recruiting RdRP enzymes to amplify siRNA pools, and perhaps by directly stabilizing sliced RNAs, poly(UG) tails could promote long-term gene silencing known to occur in *C. elegans*^85-88^. Regardless, identification of the natural RNA targets of MUT-2 should provide a powerful entree into the breadth and biological roles of poly(UG) polymerases and poly(UG) tails.

## MATERIALS AND METHODS

### Plasmid Construction

To enable overexpression of rNTases as MS2 coat protein fusions in *S. cerevisiae*, the MAP72 MS2 cassette vector was constructed. YEplac 181 (*LEU2* 2*μ*)^89^ was digested with *Hind*III and *Xho*I. Then each portion of the MS2 cassette was subcloned with unique restrictions sites, resulting in the following insert: *S. cerevisiae TEF1* promoter, MS2 coat protein, a multiple cloning site to insert the rNTase to test (consisting of *Bam*HI, *Xma*I/*Sma*I, *Not*I, *Xba*I, *Pst*I, and *Kpn*I sites), SV40 nuclear localization signal, an RGS(H_6_) sequence to verify rNTase expression by Western blotting, and *S. cerevisiae ADH1* terminator sequence.

Each rNTase tested was cloned into MAP72 by amplifying the genes indicated in Table S1 using the primers listed. All inserts were sequenced to confirm identity and lack of mutations. Site-directed mutations were made using standard methods with oligomers corresponding to the mutated sequences.

The tRNA reporter was constructed using a tRNA^His^ expression cassette, MAB812A^90^. tRNA^His^ sequence was removed by digestion with *Xho*I and *Bgl*II. Then DNA corresponding to the tRNA reporter sequence was inserted by annealing overlapping oligomers to construct both strands of the DNA sequence. The tRNA reporter is an *S. cerevisiae* tRNA^Ser(AGA)^ altered to contain an MS2 stem loop sequence (underlined) in place of the endogenous tRNA^Ser(AGA)^ variable arm (5’ - GGCAACTTGGCCGAGTGGTTAAGGCGAAAGATTAGAAATCTTT<u>ACATGAGGATCACCCATGT</u>CGC AGGTTCGAGTCCTGCAGTTGTCG-3’).

A *CCA1* cassette vector was constructed using YCplac 111 (*LEU2 CEN*)^89^ in order to express *CCA1*, SPAC1093.04, or SPCC645.10 with the same promoter and C-terminal epitope tag [RGS(H)_6_]. BY4741 yeast genomic DNA was used as a template to generate an amplicon consisting of *LEU2 CEN* vector sequence at the 5’ end, the *CCA1* promoter sequence, and a 3’ terminal sequence corresponding to the multiple cloning site of MAP72 using 5’ -GAAACAGCTATGACCATGATTACGCCAAGCTTACTAGTAGCTACTTCAGGGACAAGCAAC-3’, and 5’ - ACCCTGCAGTCTAGAAGGCGGCCGCGTGGATCCACACAAAAAAAGCCCTTATAACCCACG-3’. MAP72 was used as a template to generate an amplicon consisting of the multiple cloning site, RGS(H_6_) sequence, *ADH1* terminator sequence of MAP72, and *LEU2 CEN* vector sequence at the 3’ end using 5’ -GGATCCACGCGGCCGCCTTCTAGACTGCAGGGTACCAGAGGTTCTCACCACCACCACCAC-3’ and 5’ - CCAGTCACGACGTTGTAAAACGACGGCCAGTGAATTCCTCGAGCGGTAGAGGTGTGGTCA-3’. These two amplicons were combined with *LEU2 CEN* vector (YCplac111) linearized with *Pst*I/*Sac*I and assembled by Gibson cloning^91^. The *CCA1* cassette sequence was confirmed by Sanger sequencing. *CCA1*, SPAC1093.04, or SPCC645.10 sequences were subcloned from their respective MAP72-based constructs into the *CCA1* cassette for expression in *cca1-1* yeast.

To construct the MAP136 MUT-2 oocyte expression vector (pCS2 3HA MS2-MUT-2 WT), MUT-2a was PCR-amplified from its MAP72-based vector using 5’ -CTACCATGGATGGCTTCTAACTTTACTCAGTTCGTTCTCGTCGAC-3’ and 5’ -ACTCTCGAGTTAGTGGTGGTGGTGGTGGTGAGAACCTCTGGTACCCTGCAGTACAAATGA-3’ and then cloned into the *Nco*I/*Xho*I site of pCS2 3HA MS2. MUT-2 DNA sequence was verified prior to oocyte injections.

### Yeast Growth

BY4741 yeast were co-transformed using standard methods^92^ with a plasmid expressing the reporter RNA and a plasmid expressing the rNTase of interest, or vector controls, and selected on synthetic yeast medium lacking uracil and leucine (SD-Ura-Leu). Cultures were inoculated with single colonies, grown to saturation, and then diluted to 0.1 OD_600_/mL and grown to log phase (0.8-1 OD_600_/mL). Cells were spun down in pellets of 25 OD_600_ (approximately 5 x 10^8^ cells) and stored at −80°C until RNA extraction or protein expression analysis. We performed Western blotting with mouse anti-RGS-His Antibody (1:2500 dilution, 5PRIME/Qiagen). Only those samples with clear expression of the rNTase fusion protein were analyzed by high-throughput sequencing.

*cca1-1* yeast were co-transformed with vectors as listed in Figs. 3 and S3 using standard methods^92^, and selected on SD-Ura-Leu plates at room temperature. Colonies were selected and grown to saturation in SD-Ura-Leu liquid media. Cultures were diluted to 0.5 OD/mL followed by three 10-fold serial dilutions, spotted on SD-Ura-Leu plates, and incubated at room temperature (23°C) for 4 days or 37°C for 3 days.

### RNA Extraction

RNA was extracted from 25 OD of yeast corresponding to each sample by modification of a previously described method^93^. To each sample, 0.5 g of 0.5 mm acid washed beads (Sigma-Aldrich), 0.5 mL of RNA ISO buffer (500 mM NaCl, 200 mM Tris-Cl pH 7.5, 10 mM EDTA, 1 % SDS) and 0.5 mL of phenol-chloroform-isoamyl alcohol pH 6.7 (PCA, Fisher Scientific) was added. Samples were lysed with 10 cycles that each consisted of vortexing for 20 seconds and incubation on ice for 30 seconds. 1.5 volumes (relative to starting amount of ISO Buffer) of RNA ISO Buffer and of PCA were added, and samples were centrifuged at 4°C to separate phases. The aqueous layer was transferred to a pre-spun phase-lock gel (heavy) tube (5PRIME/Quantabio); an equal volume of PCA was added and mixed prior to centrifugation at room temperature to separate phases. The aqueous layer was transferred to 2 new tubes for ethanol precipitation with 2 volumes of 100% ethanol followed incubation at −80°C for 1 hour to overnight. Precipitated RNA was pelleted by centrifugation at 4°C. Each pellet was dissolved in 25 μL nuclease-free water and combined into 1 tube per sample. Co-purifying DNA was digested with 20 U of Turbo DNase (Invitrogen) at 37°C for 4 hours, and RNA was cleaned up with the GeneJET RNA Purification Kit (Thermo Scientific), and eluted with 50 μL of DEPC-treated water.

### RT-PCR Experiments

RT-PCR experiments to detect A tails or U tails on an RNase P RNA reporter (see Fig. S1) were performed by using a tail-specific reverse transcription step with 5 pmol of a T_33_ or A_33_ DNA primer and 100 ng of total RNA using ImProm-II Reverse Transcriptase (Promega Corporation). Then the resulting reactions were PCR-amplified using reporter-specific primers (5’ - TCGAGCCCGGGCAGCTTGCATGC-3’ and 5’ - GGGAATTCCGATCCTCTAGAGTC-3’). If a tail was added to the RNase P RNA reporter, then the RT reaction would produce cDNA, and the PCR would result in an amplicon.

RT-PCR experiments to detect tails added to the tRNA reporter were performed as described with the RNase P RNA reporter but with the following modifications. PCR amplification was performed with a forward primer specific to the 5’ end of the tRNA (5’ -GGCAACTTGGCCGAGTGGTTAAGG-3’) and a reverse primer specific to the 3’ end of the tRNA with an A tail or U tail, respectively: 5’ - AAAAAAAAAAAAAAAAAAAAAAAAAAAAAAAAATGGCGACAACTGC-3’ or 5’ - TTTTTTTTTTTTTTTTTTTTTTTTTTTTTTTTTTGGCGACAACTGC-3’. If a tail was added to the tRNA reporter, then the RT reaction would produce cDNA, and the tail-specific PCR would result in an amplicon.

### TRAID-Seq Library Preparation

Total RNA (100 ng) was ligated with 20 pmol of a 5’ adenylated primer containing a 7-nucleotide random DNA sequence (random heptamer), Illumina TruSeq adapter sequence and a 3’ dideoxycytidine (5’ - A(pp) NNNNNNN TGGAATTCTCGGGTGCCAAGG ddC-3’) using 200 U of T4 RNA ligase 2, truncated KQ (New England BioLabs) in a 20μL reaction with 16°C overnight incubation. This ligation added the random heptamer and Illumina TruSeq adapter sequence to the 3’ end of the RNAs in the sample.

Half of the ligation reaction (10 μL) was reverse transcribed using 5 pmol of Illumina RNA RT primer (5’ -GCCTTGGCACCCGAGAATTCCA-3’) and ImProm-II Reverse Transcriptase (Promega Corporation) with 1.5 mM MgCl_2_ and 0.5 mM dNTPs, according to manufacturer’s instructions.

Samples were then PCR-amplified with a forward primer consisting of Illumina-specific sequences and sequence (underlined), specific to the tRNA reporter (5’ - AATGATACGGCGACCACCGAGATCTACACGTTCAGAGTTCTACAGTCCGA<u>CGATCGAGGATCACCCATGTCGCAG</u>-3’) and a reverse Illumina RNA PCR Primer with various indices used for multiplexing, using GoTaq Green PCR Master Mix (Promega Corporation). PCR products were run on an 8% polyacrylamide 8M urea gel and gel extracted. Resulting samples for each sequencing run were combined in equimolar amounts and run on an Illumina HiSeq2000 or HiSeq2500 (2×50 bp or 2×100 bp), to produce approximately 1 x 10^6^ reads per sample.

Experiments with the RNase P RNA reporter were performed essentially as described above but with a few modifications. For TRAID-Seq, the 5’ primer used for PCR amplification was specific for the RNase P RNA reporter (5’ - AATGATACGGCGACCACCGAGATCTACACGTTCAGAGTTCTACAGTCCGACGATCGTCTGCAGGTCGACTCTAGAAA-3’).

### TRAID-Seq Data Analysis

Reads resulting from sequencing of TRAID-Seq samples were analyzed using a group of Python scripts that we call the “puppyTails” program. Briefly, puppyTails identifies sequences corresponding to the tRNA reporter, CCA end of the tRNA, and added tail in read 1. In read 2, the program identifies the random heptamer sequence, added tail sequence, and, if read length allows, the CCA end and tRNA reporter sequence. Reads were collapsed into unique ligation events using the random heptamer and then compared to identify and remove sequences resulting from PCR amplification (PCR duplicates). The number of unique times that each tail sequence is observed is counted. Tail sequences are sorted by length to calculate the nucleotide composition at each tail length and the number of tails per million heptamers (TPMH) measured for each tail length; these data are plotted as tail-o-grams (for example, Fig. 1e-g). A subsequent Perl script was used to calculate the overall nucleotide composition of tails added by a given rNTase, accounting for the number of times that a tail sequence was observed (for example, Fig. 2a-c).

### Computational Analyses of Sequence Motifs

To analyze tail sequences, a general feature screening with a random forest application^94^ was performed at the replicate level. We first quantified the number occurrences of all oligonucleotides (k=1, 2, 3, 4) within each tail sequence and utilized the resulting set of 340 features, as well as the length of the tail. The variable importance, defined as the percent mean decrease in accuracy (with 500 trees, 113 candidate variables at each split, minimum node size of 5), were estimated for all features. We define the selected features those whose importance measures are greater than 4% across replicates. We fitted a Poisson regression model in which the response variable was tail sequence counts.

#### Tails added by S. cerevisiae Cca1, S. pombe SPAC1093.04, and predicted CCA-adding enzymes

The above selected features were used as covariates. P-values from individual replicates, calculated from one-sided Wald’s test, were aggregated using Fisher’s (n<4) or Wilkinson’s (n>=4) method, followed by multiplicity correction with the Bonferroni procedure. This process identified oligonucleotides that differ between *S. cerevisiae* Cca1 and *S. pombe* SPAC1093.04 at level 0.05.

#### Tails added by C. elegans MUT-2

We evaluated the impacts of 16 dinucleotides by formally testing for their effects by a comparison of a null model without each dinucleotide and the alternative model deduced from random forest filtered set of features plus other dinucleotides. This procedure identified UG and GU as the most significant dinucleotides.

### In vitro Transcription

pCS2 3HA MS2-MUT-2 (MAP136) was linearized with *Sac*II, and 3 μg of linearized plasmid was transcribed with Ampliscribe SP6 High Yield Transcription Kit (Epicentre), according to manufacturer’s instructions. pLGMS2-luc (RNA with three MS2-binding sites)^16,95^ was linearized with *Bgl*II, and 1 μg of linearized plasmid was transcribed with T7 Flash In Vitro Transcription Kit (Epicentre), according to manufacturer’s instructions. Transcription reactions included m^7^G(5’)ppp(5’)G RNA Cap Structure Analog (New England Biolabs).

### Tethered Function Assays and Oocyte RNA Extraction

*Xenopus laevis* oocyte manipulations and injections were performed as in previous studies^16,95,96^.

Tethered function assays were conducted essentially as previously described^22^. Briefly, Stage VI oocytes were injected with 50 nL of 600 ng/µL capped mRNA encoding MS2-HA-MUT-2 protein. After 6 hours, the same oocytes were injected with 50 nL of 3 ng/µL pLGMS2-luc reporter mRNA. After 16 hours, oocytes were collected, lysed, and assayed. Three oocytes were used to confirm protein expression. Total RNA was extracted from oocytes using TRI reagent (Sigma-Aldrich), as described previously^22^, then treated with 8 U of Turbo DNase (Invitrogen) at 37°C for 1 hour, and cleaned up with the GeneJET RNA Purification Kit (Thermo Scientific).

### Oocyte RNA Analysis and Tail Sequencing

Oocyte total RNA (100 ng) was ligated with 20 pmol of the 5’ adenylated primer as described above. This ligation added the random heptamer sequence and a known sequence to the 3’ ends of RNAs in the sample for tail sequence-independent analyses. Half of the ligation reaction (10 μL) was reverse transcribed as described above.

Samples were PCR-amplified with a forward primer specific to the RNA reporter (5’ - CTCTGCAGTCGATAAAGAAAACATGAG-3’) and a reverse primer specific to the known sequence added to the 3’ end of the RNA (5’ - GCCTTGGCACCCGAGAATTCCA-3’), using GoTaq Green PCR Master Mix (Promega Corporation). PCR products were run on a 1.5% agarose gel, and purified with the GeneJET Gel Extraction Kit (Thermo Scientific). Non-templated A overhangs were added by treating the purified PCR products with 10 U of TaqPlus Precision Polymerase Mixture (Agilent Genomics) in TaqPlus Precision buffer supplemented with 0.2 mM dATP at 70°C for 30 minutes. The PCR products were then subjected to cloning with the TOPO TA Cloning Kit for Subcloning (ThermoFisher Scientific) as follows: 6% of the A addition reaction volume (2.4 μL) was combined with 0.6 μL of Salt Solution and 0.7 μL of TOPO Vector and incubated at room temperature for 30 minutes. Reactions were diluted 1 in 4 with water, transformed into DH5α competent cells, and selected on LB agar with 100 μg/mL ampicillin and 75 μg/mL X-Gal for blue/white screening. White colonies were selected, plasmids were extracted, and inserts were sequenced to identify tails added to the reporter. All reporter sequences with added tails are reported in Fig. 5f.

## ACKNOWLEDGEMENTS

We are very grateful to the Wickens and Kimble labs for advice throughout the work. We are particularly grateful to Scott Kennedy for discussions concerning MUT-2 and the manuscript, and Anita Hopper and Eric Phizicky for advice on *S. pombe* CCA-adding enzymes. We thank the Kennedy and Anderson labs for discussions. We thank the University of Wisconsin Biotechnology Center DNA Sequencing Facility, especially Marie Adams and Michael Sussmann, for providing Illumina sequencing facilities and services. We acknowledge Michael Harte for assistance with cloning *S. pombe* rNTases. We are grateful to Laura Vanderploeg of the UW Biochemistry Media Laboratory for help with the figures. This work was supported by a Ruth Kirschstein National Research Service Award (1F32GM103130-01A1) to M.A.P. and an NIH Grant to M.W (GM50942).

## AUTHOR CONTRIBUTIONS

M.A.P. and M.W. designed experiments; M.A.P. performed the experiments and analyzed data unless otherwise noted. D.F.P. wrote the PuppyTails program used to analyze TRAID-Seq data, including “tailograms.” F.C. performed statistical analyses of tail sequence motifs. N.B. prepared *N. crassa and C. albicans* TRAID-Seq samples. C.P.L wrote the Perl script used to determine total nucleotide incorporation. M.A.P. and M.W. wrote the paper, with contributions from all authors.

## SUPPLEMENTAL FIGURE LEGENDS

**Figure S1.**
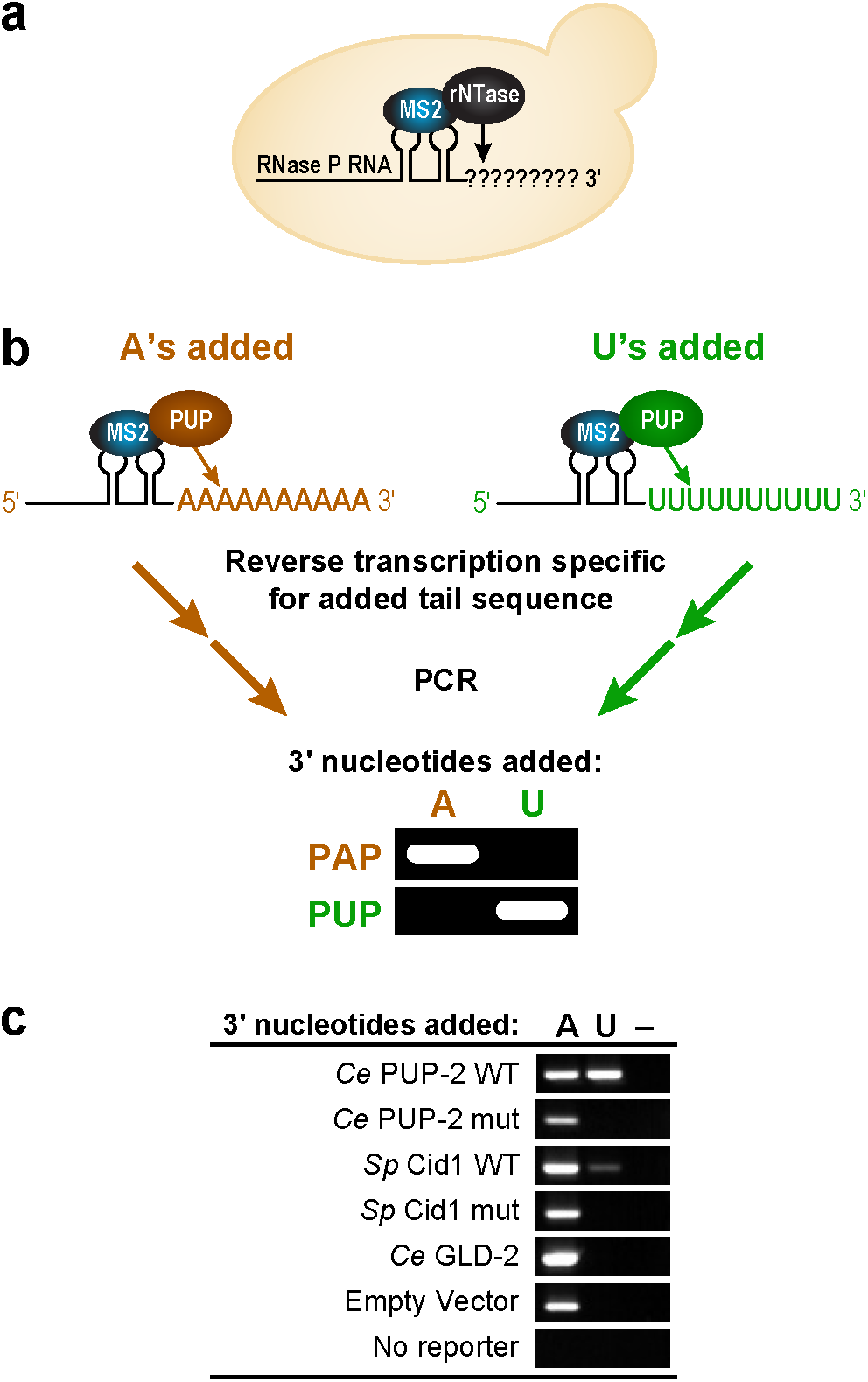
rNTase assay using RNase P-derived reporter RNA. **(a)** RNase P reporter RNA is co-expressed with an MS2 coat protein-rNTase fusion in yeast, and the tethered rNTase adds nucleotides to the 3’ end of the RNA. **(b)** Schematic of sample processing to detect A tails or U tails added to reporter RNA and example data identifying PUP or PAP activity. This approach was used to produce the RT-PCR data in Figure 1a. **(c)** RT-PCR analysis to detect A tails or U tails added to the RNase P reporter RNA by control rNTases, relative to empty vector or to a no-reporter control. Lanes marked with a dash indicate reactions performed without reverse transcriptase.

**Figure S2.**
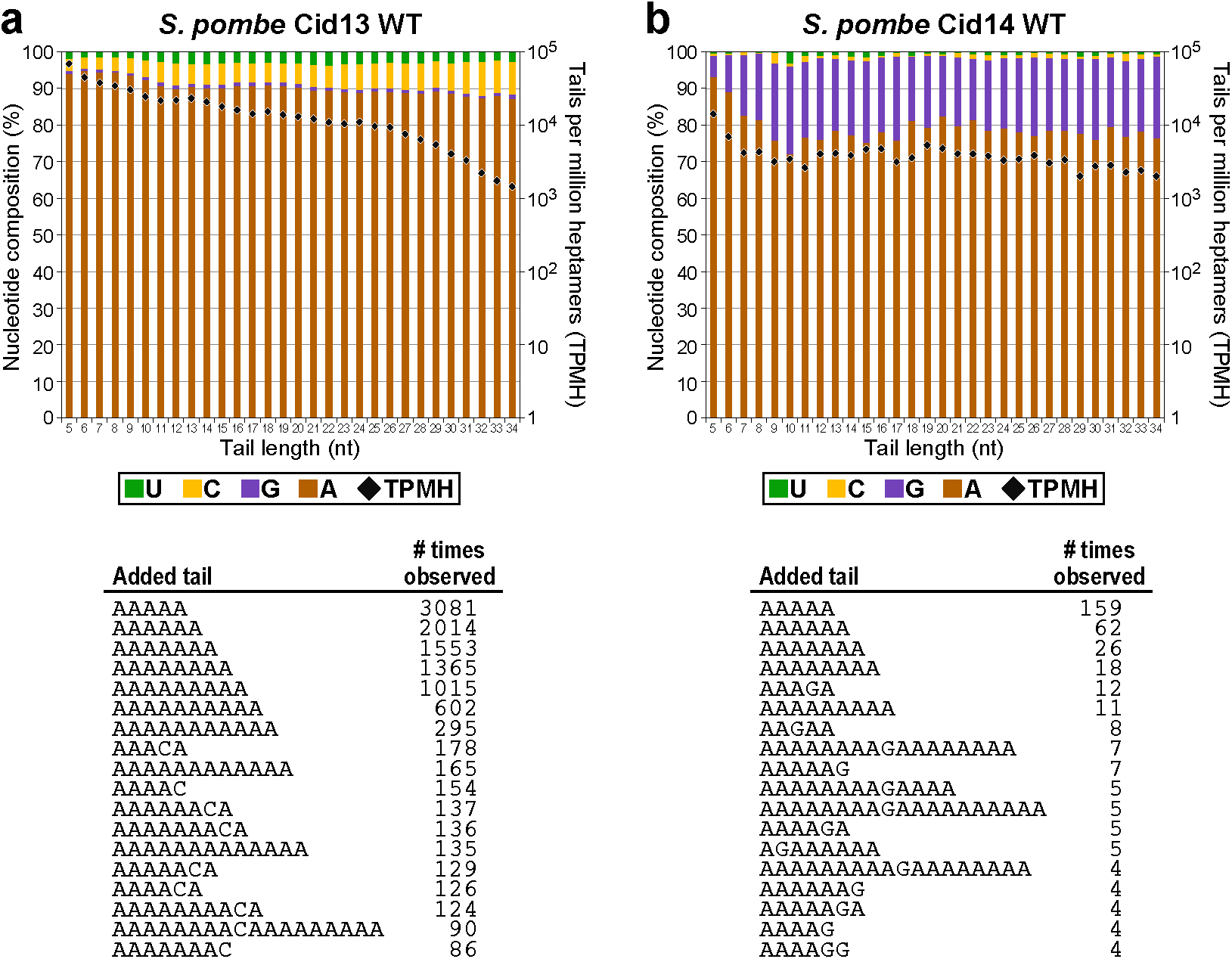
Two known *S. pombe* PAPs have secondary nucleotide preferences. **(a)** Top, tail-o-gram depicting nucleotide composition of tails added by *Sp* Cid13 and number of tails normalized to unique heptamer sequences. Bottom, representative tail sequences added to tRNA reporter. **(b)** Top, tail-o-gram depicting nucleotide composition of tails added by *Sp* Cid14 and number of tails normalized to unique heptamer sequences. Bottom, representative tail sequences added to tRNA reporter.

**Figure S3.**
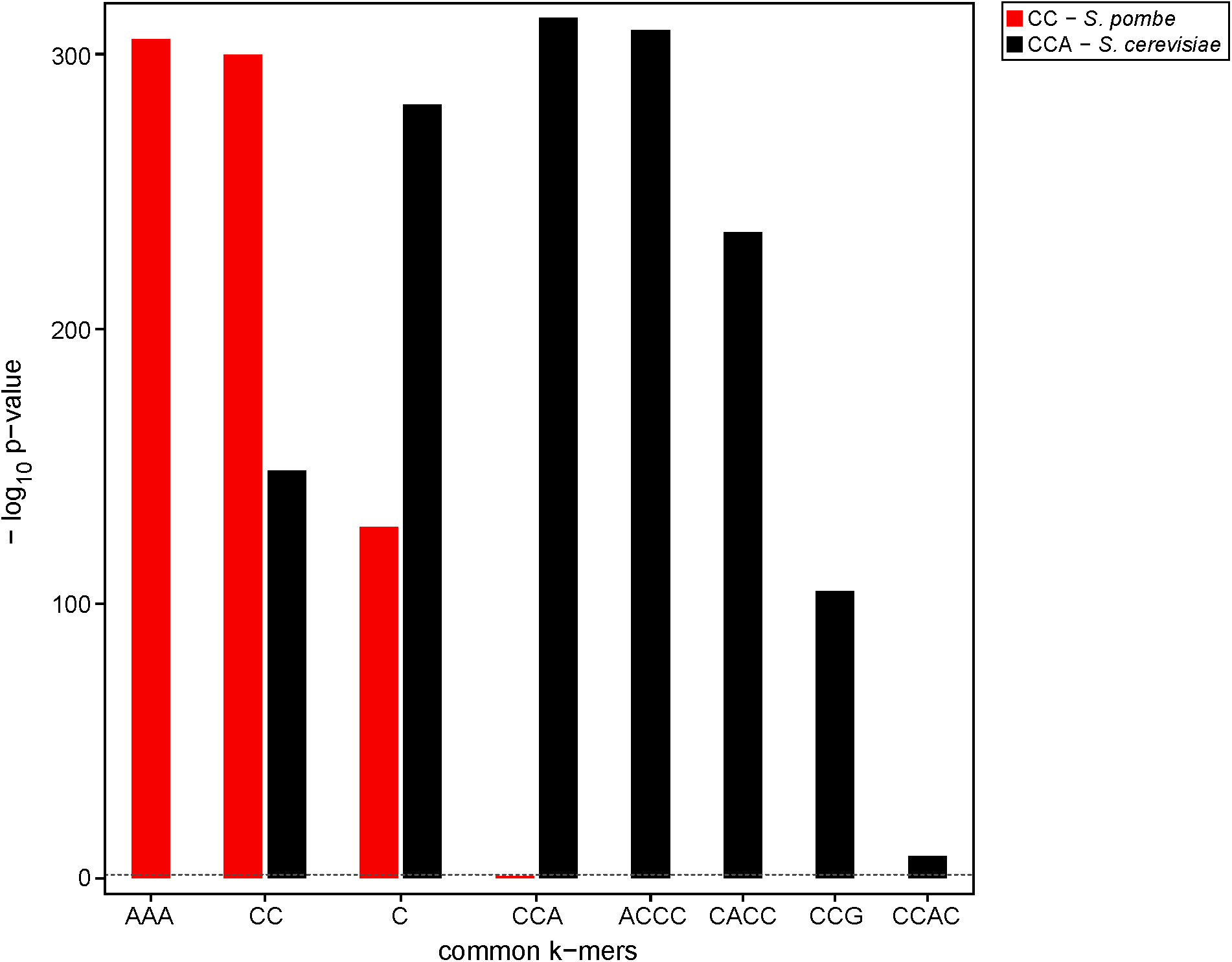
Sequence motif effect analysis of *S. pombe* and *S. cerevisiae* enzymes. Effect analysis of tails added in cells expressing *Sp* SPAC1093.04 (red) or *Sc* Cca1 (black). Displayed are the oligonucleotides with a significant effect after multiplicity correction with the Bonferroni procedure at significance level 0.05 (dashed line near baseline).

**Figure S4.**
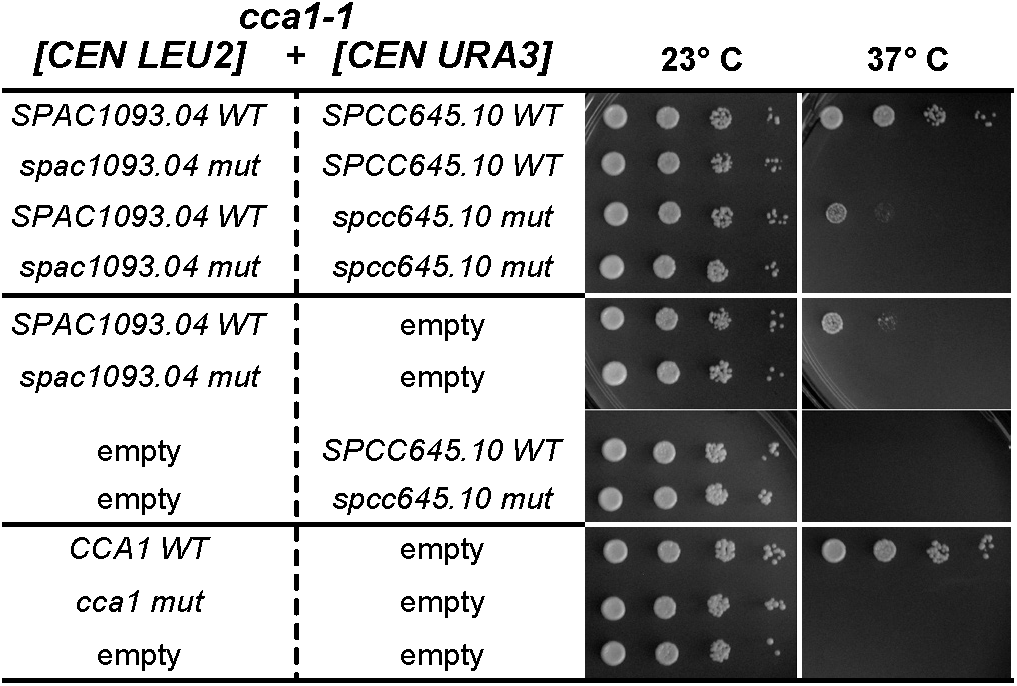
Expression of both SPAC1093.04 and SPCC645.10 rescues *cca1-1* temperature sensitivity: second biological replicate. *cca1-1* mutant strains containing *CEN* plasmids expressing indicated plasmids were serially diluted, spotted on SD-Ura-Leu media and grown at 37°C for 3 days or 23°C for 4 days.

**Figure S5.**
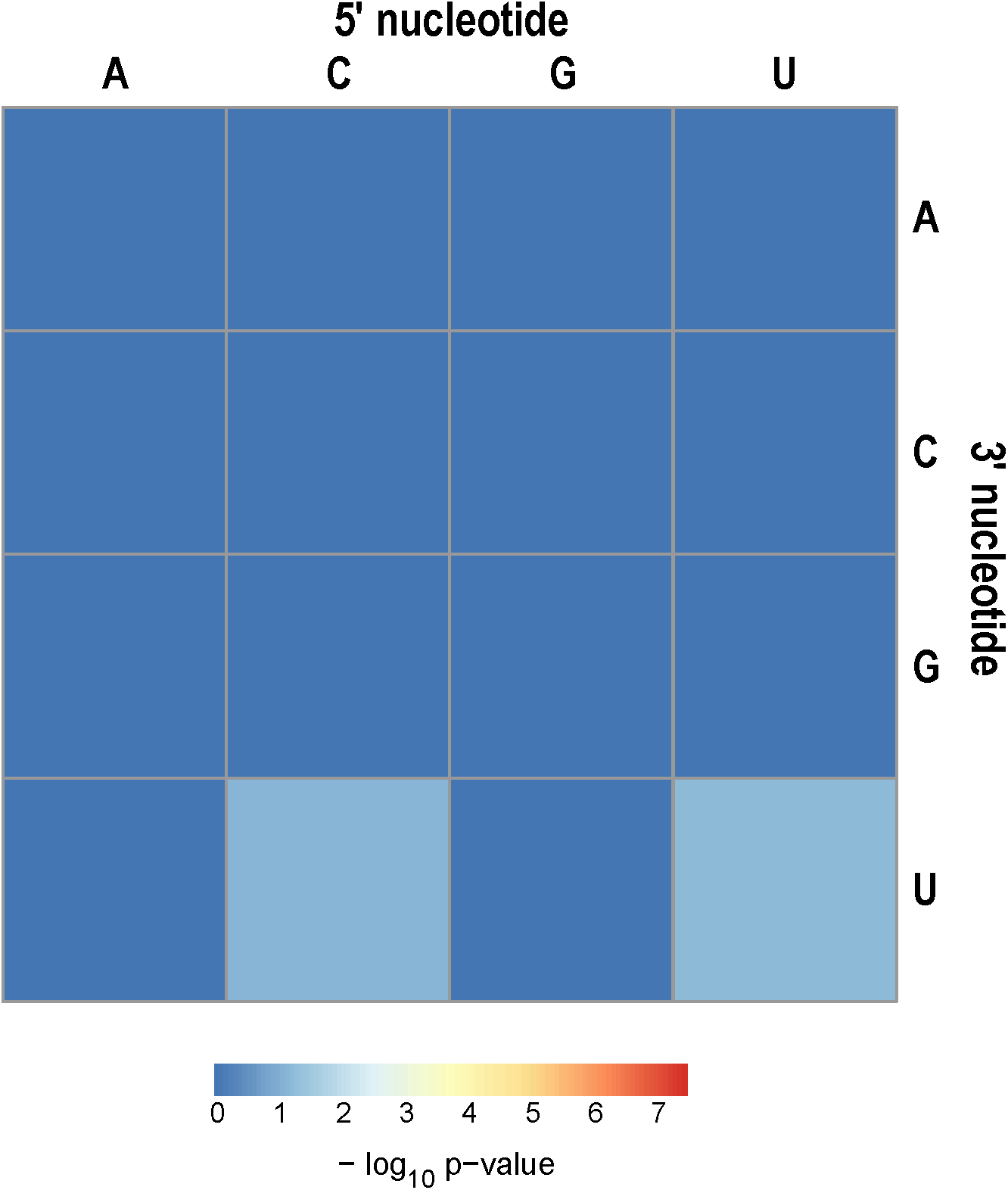
Statistical analysis of all possible dinucleotides in tails added by NPOL-1. Heatmap of p-values for all possible dinucleotides with negative logarithm (base 10) is presented. No specific dinucleotide had a statistically significant effect after multiplicity correction with the Bonferroni procedure at significance level 0.05.

**Table S1.**
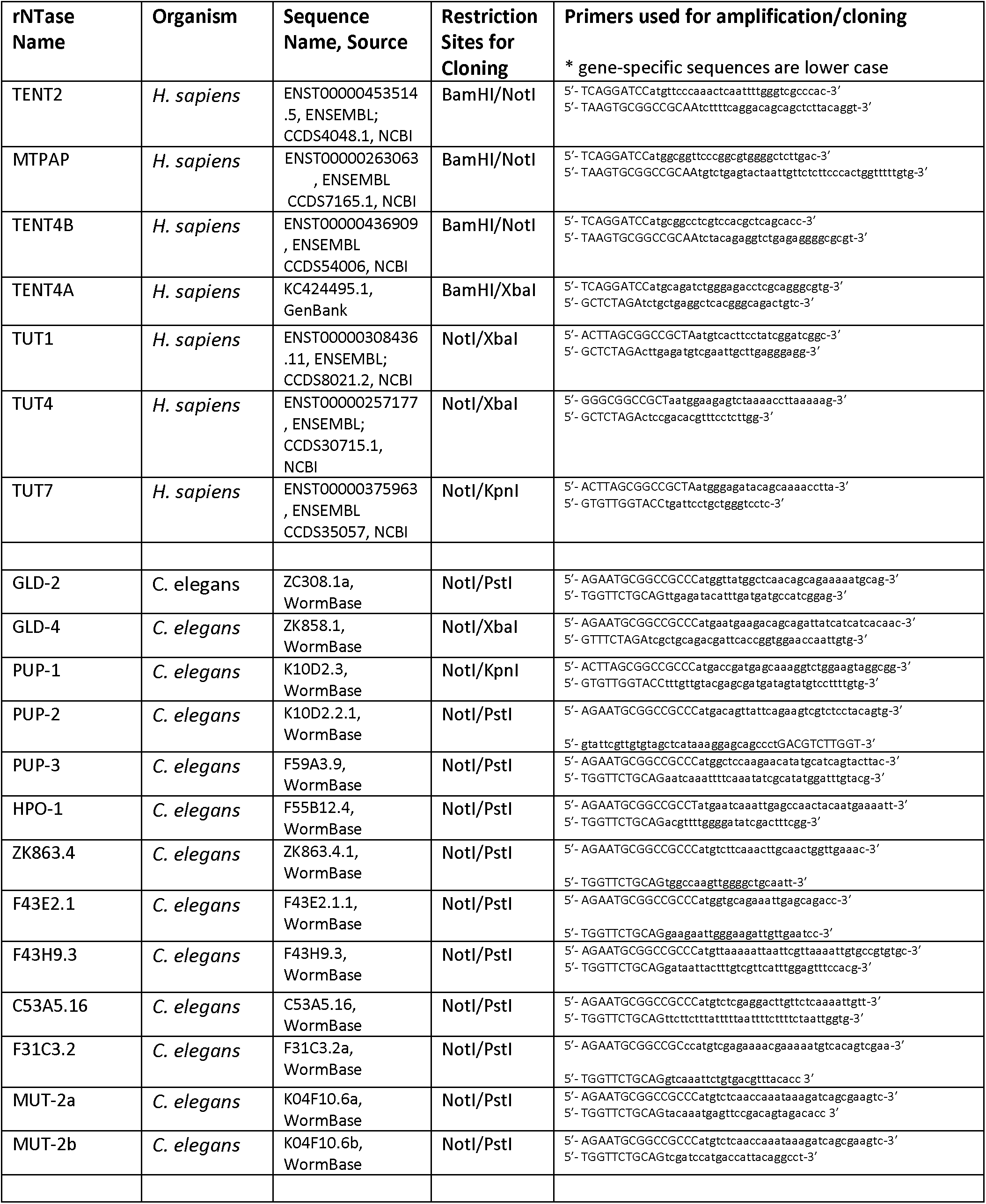
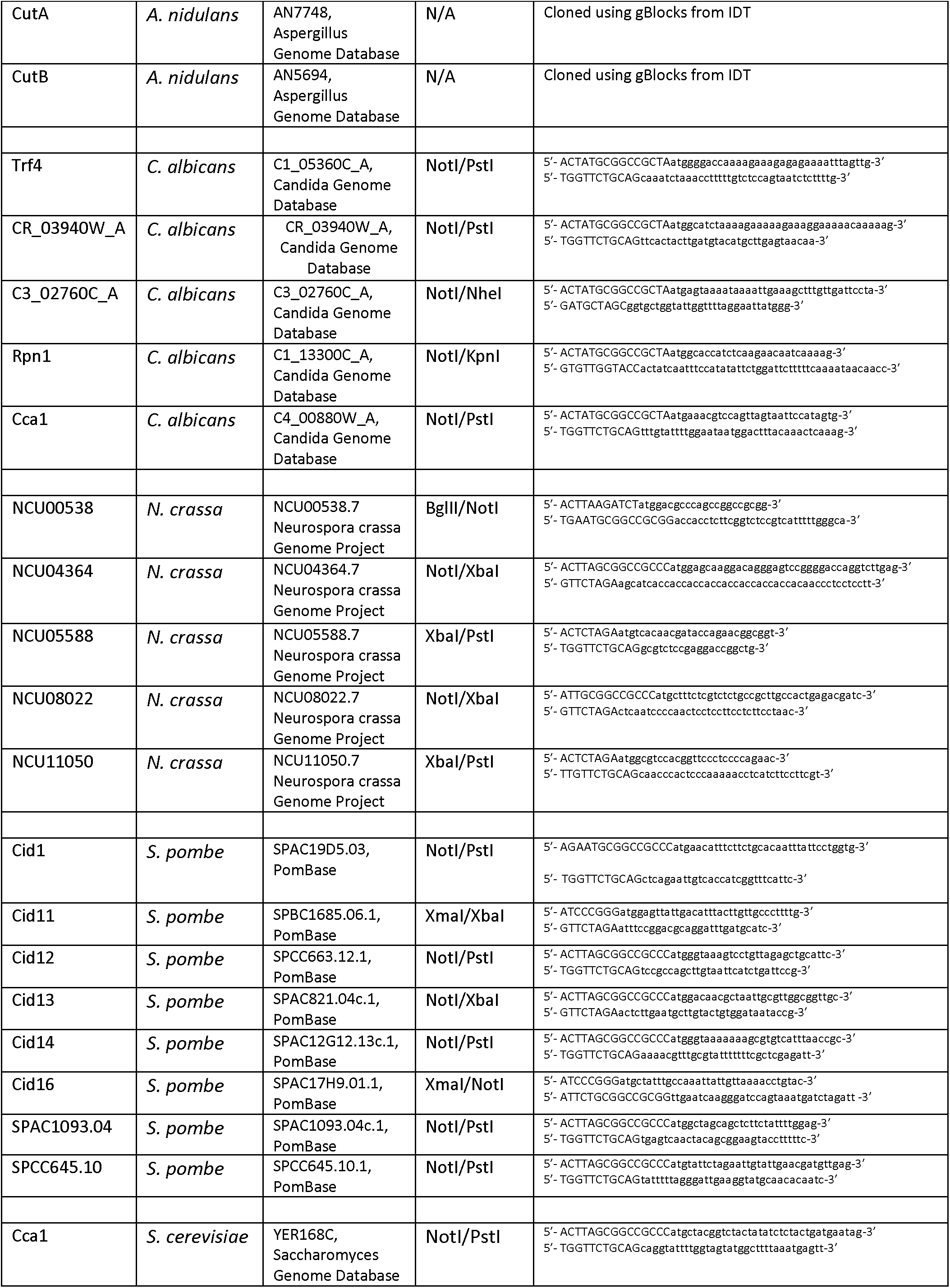
rNTase plasmid construction information

## REFERENCES

1 Blahna, M. T., Jones, M. R., Quinton, L. J., Matsuura, K. Y. & Mizgerd, J. P. Terminal uridyltransferase enzyme Zcchc11 promotes cell proliferation independent of its uridyltransferase activity. J Biol Chem 286, 42381–42389, doi:10.1074/jbc.M111.259689 (2011).

2 Chang, H. et al. Terminal Uridylyltransferases Execute Programmed Clearance of Maternal Transcriptome in Vertebrate Embryos. Mol Cell 70, 72–82 e77, doi:10.1016/j.molcel.2018.03.004 (2018).

3 Gutierrez-Vazquez, C. et al. 3’ Uridylation controls mature microRNA turnover during CD4 T-cell activation. RNA 23, 882–891, doi:10.1261/rna.060095.116 (2017).

4 Hagan, J. P., Piskounova, E. & Gregory, R. I. Lin28 recruits the TUTase Zcchc11 to inhibit let-7 maturation in mouse embryonic stem cells. Nat Struct Mol Biol 16, 1021–1025, doi:10.1038/nsmb.1676 (2009).

5 Huo, Y. et al. Widespread 3’-end uridylation in eukaryotic RNA viruses. Sci Rep 6, 25454, doi:10.1038/srep25454 (2016).

6 Jones, M. R. et al. Zcchc11 uridylates mature miRNAs to enhance neonatal IGF-1 expression, growth, and survival. PLoS Genet 8, e1003105, doi:10.1371/journal.pgen.1003105 (2012).

7 Jones, M. R. et al. Zcchc11-dependent uridylation of microRNA directs cytokine expression. Nat Cell Biol 11, 1157–1163, doi:10.1038/ncb1931 (2009).

8 Minoda, Y. et al. A novel Zinc finger protein, ZCCHC11, interacts with TIFA and modulates TLR signaling. Biochem Biophys Res Commun 344, 1023–1030, doi:10.1016/j.bbrc.2006.04.006 (2006).

9 Piskounova, E. et al. Lin28A and Lin28B Inhibit let-7 MicroRNA Biogenesis by Distinct Mechanisms. Cell 147, 1066–1079, doi:10.1016/j.cell.2011.10.039 (2011).

10 van Wolfswinkel, J. C. et al. CDE-1 affects chromosome segregation through uridylation of CSR-1-bound siRNAs. Cell 139, 135–148, doi:10.1016/j.cell.2009.09.012 (2009).

11 A, P. S. & Laishram, R. S. Nuclear Phosphatidylinositol-Phosphate Type I Kinase alpha-Coupled Star-PAP Polyadenylation Regulates Cell Invasion. Mol Cell Biol 38, doi:10.1128/MCB.00457-17 (2018).

12 Benoit, P., Papin, C., Kwak, J. E., Wickens, M. & Simonelig, M. PAP- and GLD-2-type poly(A) polymerases are required sequentially in cytoplasmic polyadenylation and oogenesis in *Drosophila*. Development 135, 1969–1979, doi:10.1242/dev.021444 (2008).

13 Burns, D. M., D’Ambrogio, A., Nottrott, S. & Richter, J. D. CPEB and two poly(A) polymerases control miR-122 stability and p53 mRNA translation. Nature 473, 105–108, doi:10.1038/nature09908 (2011).

14 Kim, K. W., Wilson, T. L. & Kimble, J. GLD-2/RNP-8 cytoplasmic poly(A) polymerase is a broad-spectrum regulator of the oogenesis program. Proc Natl Acad Sci U S A 107, 17445–17450, doi:10.1073/pnas.1012611107 (2010).

15 Kwak, J. E. et al. GLD2 poly(A) polymerase is required for long-term memory. Proc Natl Acad Sci U S A 105, 14644–14649, doi:10.1073/pnas.0803185105 (2008).

16 Kwak, J. E., Wang, L., Ballantyne, S., Kimble, J. & Wickens, M. Mammalian GLD-2 homologs are poly(A) polymerases. Proc Natl Acad Sci U S A 101, 4407–4412, doi:10.1073/pnas.0400779101 (2004).

17 Yu, C. et al. Star-PAP, a poly(A) polymerase, functions as a tumor suppressor in an orthotopic human breast cancer model. Cell Death Dis 8, e2582, doi:10.1038/cddis.2016.199 (2017).

18 Yue, D., Maizels, N. & Weiner, A. M. CCA-adding enzymes and poly(A) polymerases are all members of the same nucleotidyltransferase superfamily: characterization of the CCA-adding enzyme from the archaeal hyperthermophile Sulfolobus shibatae. RNA 2, 895–908(1996).

19 Aravind, L. & Koonin, E. V. DNA polymerase beta-like nucleotidyltransferase superfamily: identification of three new families, classification and evolutionary history. Nucleic Acids Res 27, 1609–1618(1999).

20 Martin, G. & Keller, W. RNA-specific ribonucleotidyl transferases. RNA 13, 1834–1849, doi:10.1261/rna.652807 (2007).

21 Mellman, D. L. et al. A PtdIns4,5P2-regulated nuclear poly(A) polymerase controls expression of select mRNAs. Nature 451, 1013–1017, doi:10.1038/nature06666 (2008).

22 Kwak, J. E. & Wickens, M. A family of poly(U) polymerases. RNA 13, 860–867, doi:10.1261/rna.514007 (2007).

23 Lapointe, C. P. & Wickens, M. The nucleic acid-binding domain and translational repression activity of a Xenopus terminal uridylyl transferase. J Biol Chem 288, 20723–20733, doi:10.1074/jbc.M113.455451 (2013).

24 Chang, H., Lim, J., Ha, M. & Kim, V. N. TAIL-seq: genome-wide determination of poly(A) tail length and 3’ end modifications. Mol Cell 53, 1044–1052, doi:10.1016/j.molcel.2014.02.007 (2014).

25 Gazestani, V. H., Hampton, M., Abrahante, J. E., Salavati, R. & Zimmer, S. L. circTAIL-seq, a targeted method for deep analysis of RNA 3’ tails, reveals transcript-specific differences by multiple metrics. RNA 22, 477–486, doi:10.1261/rna.054494.115 (2016).

26 Lim, J., Lee, M., Son, A., Chang, H. & Kim, V. N. mTAIL-seq reveals dynamic poly(A) tail regulation in oocyte-to-embryo development. Genes Dev 30, 1671–1682, doi:10.1101/gad.284802.116 (2016).

27 Collins, J., Saari, B. & Anderson, P. Activation of a transposable element in the germ line but not the soma of Caenorhabditis elegans. Nature 328, 726–728, doi:10.1038/328726a0 (1987).

28 Yuan, J. Y., Finney, M., Tsung, N. & Horvitz, H. R. Tc4, a Caenorhabditis elegans transposable element with an unusual fold-back structure. Proc Natl Acad Sci U S A 88, 3334–3338(1991).

29 Collins, J. J. & Anderson, P. The Tc5 family of transposable elements in Caenorhabditis elegans. Genetics 137, 771–781(1994).

30 Tabara, H. et al. The rde-1 gene, RNA interference, and transposon silencing in C. elegans. Cell 99, 123–132 (1999).

31 Ketting, R. F. & Plasterk, R. H. A genetic link between co-suppression and RNA interference in C. elegans. Nature 404, 296–298, doi:10.1038/35005113 (2000).

32 Dernburg, A. F., Zalevsky, J., Colaiacovo, M. P. & Villeneuve, A. M. Transgene-mediated cosuppression in the C. elegans germ line. Genes Dev 14, 1578–1583 (2000).

33 Robert, V. J., Sijen, T., van Wolfswinkel, J. & Plasterk, R. H. Chromatin and RNAi factors protect the *C. elegans* germline against repetitive sequences. Genes Dev 19, 782–787, doi:10.1101/gad.332305 (2005).

34 Lee, H. C. et al. C. elegans piRNAs mediate the genome-wide surveillance of germline transcripts. Cell 150, 78–87, doi:10.1016/j.cell.2012.06.016 (2012).

35 Chen, C. C. et al. A member of the polymerase beta nucleotidyltransferase superfamily is required for RNA interference in *C. elegans*. Curr Biol 15, 378–383, doi:10.1016/j.cub.2005.01.009 (2005).

36 Lee, R. C., Hammell, C. M. & Ambros, V. Interacting endogenous and exogenous RNAi pathways in *Caenorhabditis elegans*. RNA 12, 589–597, doi:10.1261/rna.2231506 (2006).

37 Gu, W. et al. Distinct argonaute-mediated 22G-RNA pathways direct genome surveillance in the *C. elegans* germline. Mol Cell 36, 231–244, doi:10.1016/j.molcel.2009.09.020 (2009).

38 Jose, A. M., Garcia, G. A. & Hunter, C. P. Two classes of silencing RNAs move between *Caenorhabditis elegans* tissues. Nat Struct Mol Biol 18, 1184–1188, doi:10.1038/nsmb.2134 (2011).

39 Zhang, C. et al. mut-16 and other mutator class genes modulate 22G and 26G siRNA pathways in *Caenorhabditis elegans*. Proc Natl Acad Sci U S A 108, 1201–1208, doi:10.1073/pnas.1018695108 (2011).

40 Coller, J. M., Gray, N. K. & Wickens, M. P. mRNA stabilization by poly(A) binding protein is independent of poly(A) and requires translation. Genes Dev 12, 3226–3235 (1998).

41 SenGupta, D. J. et al. A three-hybrid system to detect RNA-protein interactions in vivo. Proc Natl Acad Sci U S A 93, 8496–8501 (1996).

42 Stumpf, C. R., Opperman, L. & Wickens, M. Chapter 14. Analysis of RNA-protein interactions using a yeast three-hybrid system. Methods Enzymol 449, 295–315, doi:10.1016/S0076-6879(08)02414-2 (2008).

43 Rissland, O. S., Mikulasova, A. & Norbury, C. J. Efficient RNA polyuridylation by noncanonical poly(A) polymerases. Mol Cell Biol 27, 3612–3624, doi:10.1128/MCB.02209-06 (2007).

44 Wang, L., Eckmann, C. R., Kadyk, L. C., Wickens, M. & Kimble, J. A regulatory cytoplasmic poly(A) polymerase in *Caenorhabditis elegans*. Nature 419, 312–316, doi:10.1038/nature01039 (2002).

45 Read, R. L., Martinho, R. G., Wang, S. W., Carr, A. M. & Norbury, C. J. Cytoplasmic poly(A) polymerases mediate cellular responses to S phase arrest. Proc Natl Acad Sci U S A 99, 12079–12084, doi:10.1073/pnas.192467799 (2002).

46 Saitoh, S. et al. Cid13 is a cytoplasmic poly(A) polymerase that regulates ribonucleotide reductase mRNA. Cell 109, 563–573 (2002).

47 Kadaba, S. et al. Nuclear surveillance and degradation of hypomodified initiator tRNAMet in *S. cerevisiae*. Genes Dev 18, 1227–1240, doi:10.1101/gad.1183804 (2004).

48 Tomecki, R., Dmochowska, A., Gewartowski, K., Dziembowski, A. & Stepien, P. P. Identification of a novel human nuclear-encoded mitochondrial poly(A) polymerase. Nucleic Acids Res 32, 6001–6014, doi:10.1093/nar/gkh923 (2004).

49 Haracska, L., Johnson, R. E., Prakash, L. & Prakash, S. Trf4 and Trf5 proteins of Saccharomyces cerevisiae exhibit poly(A) RNA polymerase activity but no DNA polymerase activity. Mol Cell Biol 25, 10183–10189, doi:10.1128/MCB.25.22.10183-10189.2005 (2005).

50 Kadaba, S., Wang, X. & Anderson, J. T. Nuclear RNA surveillance in *Saccharomyces cerevisiae*: Trf4p-dependent polyadenylation of nascent hypomethylated tRNA and an aberrant form of 5S rRNA. RNA 12, 508–521, doi:10.1261/rna.2305406 (2006).

51 Trippe, R. et al. Identification, cloning, and functional analysis of the human U6 snRNA-specific terminal uridylyl transferase. RNA 12, 1494–1504, doi:10.1261/rna.87706 (2006).

52 Schmid, M., Kuchler, B. & Eckmann, C. R. Two conserved regulatory cytoplasmic poly(A) polymerases, GLD-4 and GLD-2, regulate meiotic progression in C. elegans. Genes Dev 23, 824–836, doi:10.1101/gad.494009 (2009).

53 Rammelt, C., Bilen, B., Zavolan, M. & Keller, W. PAPD5, a noncanonical poly(A) polymerase with an unusual RNA-binding motif. RNA 17, 1737–1746, doi:10.1261/rna.2787011 (2011).

54 Phizicky, E. M. & Hopper, A. K. tRNA processing, modification, and subcellular dynamics: past, present, and future. RNA 21, 483–485, doi:10.1261/rna.049932.115 (2015).

55 Tomita, K. & Weiner, A. M. Collaboration between CC- and A-adding enzymes to build and repair the 3’-terminal CCA of tRNA in Aquifex aeolicus. Science 294, 1334–1336, doi:10.1126/science.1063816 (2001).

56 Tomita, K. & Weiner, A. M. Closely related CC- and A-adding enzymes collaborate to construct and repair the 3’-terminal CCA of tRNA in Synechocystis sp. and Deinococcus radiodurans. J Biol Chem 277, 48192–48198, doi:10.1074/jbc.M207527200 (2002).

57 Aebi, M. et al. Isolation of a temperature-sensitive mutant with an altered tRNA nucleotidyltransferase and cloning of the gene encoding tRNA nucleotidyltransferase in the yeast Saccharomyces cerevisiae. J Biol Chem 265, 16216–16220 (1990).

58 Chen, J. Y., Kirchner, G., Aebi, M. & Martin, N. C. Purification and properties of yeast ATP (CTP):tRNA nucleotidyltransferase from wild type and overproducing cells. J Biol Chem 265, 16221–16224 (1990).

59 Kim, D. U. et al. Analysis of a genome-wide set of gene deletions in the fission yeast Schizosaccharomyces pombe. Nat Biotechnol 28, 617–623, doi:10.1038/nbt.1628 (2010).

60 Hayles, J. et al. A genome-wide resource of cell cycle and cell shape genes of fission yeast. Open Biol 3, 130053, doi:10.1098/rsob.130053 (2013).

61 Nick McElhinny, S. A. et al. Abundant ribonucleotide incorporation into DNA by yeast replicative polymerases. Proc Natl Acad Sci U S A 107, 4949–4954, doi:10.1073/pnas.0914857107 (2010).

62 Chung, C. Z., Jo, D. H. & Heinemann, I. U. Nucleotide specificity of the human terminal nucleotidyltransferase Gld2 (TUT2). RNA 22, 1239–1249, doi:10.1261/rna.056077.116 (2016).

63 Munoz-Tello, P., Gabus, C. & Thore, S. Functional implications from the Cid1 poly(U) polymerase crystal structure. Structure 20, 977–986, doi:10.1016/j.str.2012.04.006 (2012).

64 Lunde, B. M., Magler, I. & Meinhart, A. Crystal structures of the Cid1 poly (U) polymerase reveal the mechanism for UTP selectivity. Nucleic Acids Res, doi:10.1093/nar/gks7402012).

65 Yates, L. A. et al. Structural basis for the activity of a cytoplasmic RNA terminal uridylyl transferase. Nat Struct Mol Biol 19, 782–787, doi:10.1038/nsmb.2329 (2012).

66 Munoz-Tello, P., Gabus, C. & Thore, S. A critical switch in the enzymatic properties of the Cid1 protein deciphered from its product-bound crystal structure. Nucleic Acids Res 42, 3372–3380, doi:10.1093/nar/gkt1278 (2014).

67 Yates, L. A. et al. Structural plasticity of Cid1 provides a basis for its distributive RNA terminal uridylyl transferase activity. Nucleic Acids Res 43, 2968–2979, doi:10.1093/nar/gkv122 (2015).

68 Mellman, D. L. & Anderson, R. A. A novel gene expression pathway regulated by nuclear phosphoinositides. Adv Enzyme Regul 49, 11–28 (2009).

69 Li, W., Laishram, R. S. & Anderson, R. A. The novel poly(A) polymerase Star-PAP is a signal-regulated switch at the 3’-end of mRNAs. Adv Biol Regul 53, 64–76, doi:10.1016/j.jbior.2012.10.004 (2013).

70 Morozov, I. Y., Jones, M. G., Razak, A. A., Rigden, D. J. & Caddick, M. X. CUCU modification of mRNA promotes decapping and transcript degradation in Aspergillus nidulans. Mol Cell Biol 30, 460–469, doi:10.1128/MCB.00997-09 (2010).

71 Kobylecki, K., Kuchta, K., Dziembowski, A., Ginalski, K. & Tomecki, R. Biochemical and structural bioinformatics studies of fungal CutA nucleotidyltransferases explain their unusual specificity toward CTP and increased tendency for cytidine incorporation at the 3’-terminal positions of synthesized tails. RNA 23, 1902–1926, doi:10.1261/rna.061010.117 (2017).

72 Morozov, I. Y. et al. mRNA 3’ tagging is induced by nonsense-mediated decay and promotes ribosome dissociation. Mol Cell Biol 32, 2585–2595, doi:10.1128/MCB.00316-12(2012).

73 Lim, J. et al. Mixed tailing by TENT4A and TENT4B shields mRNA from rapid deadenylation. Science, doi:10.1126/science.aam5794>(2018).

74 Boule, J. B., Rougeon, F. & Papanicolaou, C. Terminal deoxynucleotidyl transferase indiscriminately incorporates ribonucleotides and deoxyribonucleotides. J Biol Chem 276, 31388–31393, doi:10.1074/jbc.M105272200 (2001).

75 Seth, M., Thurlow, D. L. & Hou, Y. M. Poly(C) synthesis by class I and class II CCA-adding enzymes. Biochemistry 41, 4521–4532 (2002).

76 Cho, H. D., Verlinde, C. L. & Weiner, A. M. Reengineering CCA-adding enzymes to function as (U,G)- or dCdCdA-adding enzymes or poly(C,A) and poly(U,G) polymerases. Proc Natl Acad Sci U S A 104, 54–59, doi:10.1073/pnas.0606961104 (2007).

77 Fox, C. A. & Wickens, M. Poly(A) removal during oocyte maturation: a default reaction selectively prevented by specific sequences in the 3’ UTR of certain maternal mRNAs. Genes Dev 4, 2287– 2298 (1990).

78 Yue, D., Weiner, A. M. & Maizels, N. The CCA-adding enzyme has a single active site. J Biol Chem 273, 29693–29700 (1998).

79 Augustin, M. A. et al. Crystal structure of the human CCA-adding enzyme: insights into template-independent polymerization. J Mol Biol 328, 985–994 (2003).

80 Cho, H. D. & Weiner, A. M. A single catalytically active subunit in the multimeric Sulfolobus shibatae CCA-adding enzyme can carry out all three steps of CCA addition. J Biol Chem 279, 40130–40136, doi:10.1074/jbc.M405518200 (2004).

81 Xiong, Y. & Steitz, T. A. Mechanism of transfer RNA maturation by CCA-adding enzyme without using an oligonucleotide template. Nature 430, 640–645, doi:10.1038/nature02711 (2004).

82 Cho, H. D., Verlinde, C. L. & Weiner, A. M. Archaeal CCA-adding enzymes: central role of a highly conserved beta-turn motif in RNA polymerization without translocation. J Biol Chem 280, 9555–9566, doi:10.1074/jbc.M412594200 (2005).

83 Wilusz, J. E., Whipple, J. M., Phizicky, E. M. & Sharp, P. A. tRNAs marked with CCACCA are targeted for degradation. Science 334, 817–821, doi:10.1126/science.1213671 (2011).

84 Wende, S., Bonin, S., Gotze, O., Betat, H. & Morl, M. The identity of the discriminator base has an impact on CCA addition. Nucleic Acids Res 43, 5617–5629, doi:10.1093/nar/gkv471 (2015).

85 Vastenhouw, N. L. et al. Gene expression: long-term gene silencing by RNAi. Nature 442, 882, doi:10.1038/442882a (2006).

86 Buckley, B. A. et al. A nuclear Argonaute promotes multigenerational epigenetic inheritance and germline immortality. Nature 489, 447–451, doi:10.1038/nature11352 (2012).

87 Fire, A. et al. Potent and specific genetic interference by double-stranded RNA in *Caenorhabditis elegans*. Nature 391, 806–811, doi:10.1038/35888 (1998).

88 Grishok, A., Tabara, H. & Mello, C. C. Genetic requirements for inheritance of RNAi in C. elegans. Science 287, 2494–2497 (2000).

89 Gietz, R. D. & Sugino, A. New yeast-Escherichia coli shuttle vectors constructed with in vitro mutagenized yeast genes lacking six-base pair restriction sites. Gene 74, 527–534 (1988).

90 Whipple, J. M., Lane, E. A., Chernyakov, I., D’Silva, S. & Phizicky, E. M. The yeast rapid tRNA decay pathway primarily monitors the structural integrity of the acceptor and T-stems of mature tRNA. Genes Dev 25, 1173–1184, doi:10.1101/gad.2050711 (2011).

91 Gibson, D. G. et al. Enzymatic assembly of DNA molecules up to several hundred kilobases. Nat Methods 6, 343–345, doi:10.1038/nmeth.1318 (2009).

92 Sherman, F. Getting started with yeast. Methods Enzymol 350, 3–41 (2002).

93 Preston, M. A., D’Silva, S., Kon, Y. & Phizicky, E. M. tRNAHis 5-methylcytidine levels increase in response to several growth arrest conditions in Saccharomyces cerevisiae. RNA 19, 243–256, doi:10.1261/rna.035808.112 (2013).

94 Liaw, A. & Wiener, M. Classification and Regression by randomForest. R News 2, 18–22 (2002).

95 Dickson, K. S., Thompson, S. R., Gray, N. K. & Wickens, M. Poly(A) polymerase and the regulation of cytoplasmic polyadenylation. J Biol Chem 276, 41810–41816, doi:10.1074/jbc.M103030200 (2001).

96 Gray, N. K., Coller, J. M., Dickson, K. S. & Wickens, M. Multiple portions of poly(A)-binding protein stimulate translation in vivo. EMBO J 19, 4723–4733, doi:10.1093/emboj/19.17.4723 (2000).

